# An anatomically distinct dopaminergic cell population of the zona incerta as evidence for the human A13 nucleus

**DOI:** 10.64898/2026.05.11.724328

**Authors:** Violet M. Liu, Roy AM Haast, Tallulah S. Andrews, Wataru Inoue, Alaa Taha, Jason Kai, Ali R. Khan, Jonathan C. Lau

## Abstract

The zona incerta (ZI) is a functionally and molecularly diverse deep brain region increasingly recognized as an integrative hub. The rodent ZI contains the dopaminergic A13 nucleus, with studies supporting a central role in modulating sensorimotor integration, nociception, and confers resilience in models of neurodegeneration; however, to date, no clear human homologue has been identified. Here, we identified a tyrosine hydroxylase-positive subregion in the human ZI consistent with the A13, and refined its borders using *ex vivo* MRI and histology. We translate these findings to *in vivo* MRI, demonstrating the A13 is distinguishable from surrounding regions by elevated T_1_ and T_2_* values. We further revealed a principal rostral-caudal axis of molecular and structural connectivity variation in the ZI, and identified the A13 as a localized molecular specialization. Altogether, we report a distinct ZI subregion corresponding to the A13 nucleus, providing an anatomical foundation for further mechanistic and translational investigation.

## Introduction

Initially described as “a region for which nothing can be said”, our understanding of the zona incerta (ZI) has evolved since its first description by Forel in 1877^1^. Located ventral to the thalamus, the ZI is bordered by white matter tracts collectively known as the fields of Forel, which include the thalamic fasciculus (H1), lenticular fasciculus (H2), and the H field (convergence of H1 and H2)^2–4^. Growing evidence supports the central role of the ZI as an integrative hub^5–7^, due to its widespread cortical^5,8–12^ and subcortical connections^5,13–16^. These connections position the ZI to modulate diverse processes including anxiety and fear response^17–21^, novelty-seeking^22,23^, pain perception^24–28^, memory^29–31^, and sleep-wake regulation^32,33^.

Traditionally, the ZI has been parcellated into discrete subregions based on neurochemical differences, including a rostral subregion (rZI) enriched with somatostatin-positive GABAergic neurons^17,34^, and a caudal subregion (cZI) populated with glutamatergic neurons^35,36^. However, the organization of neurons may also be understood as continuous “gradients”, or transitions in molecular profiles along different spatial axes^37^. This gradient-based framework is an organizational principle widely observed throughout the brain^38^, seen along the canonical sensory-association axis of the cortex^38–41^, and within subcortical structures including the thalamus^37,42^, and the nearby subthalamic nucleus (STN)^43^. The concept of gradient-based organization can be used to describe a variety of features in the brain including neurochemical profiles^37^, structural^38^, or functional connections^44,45^, and translated across scales. Indeed, human cortico-incertal structural connectivity has been shown to follow a continuous, topographical organization along the rostral-caudal axis^10,11^.

However, macroscale connectivity alone lacks the ability to resolve subregions defined by neurochemical signatures. As there have been no cross-scale investigations bridging the connectivity-driven gradients to the underlying molecular architecture in the human ZI, neurochemically defined subregions remain unmapped. A prime example is the A13 nucleus, a dense cluster of tyrosine hydroxylase (TH)-positive neurons in the rostromedial region of the ZI (rmZI)^46,47^. The A13 is a diencephalic component of the dopaminergic system^48,49^. Functionally, this specialized cluster is postulated to modulate sensory-motor integration^50–52^, pain processing^53,54^, and appetitive behaviours^55,56^. It may also confer resilience to neurodegenerative pathology, as A13 stimulation can restore locomotion deficient in Parkinson’s disease rat models^50^. Although the identification of a TH+ region in the ZI across rodents^57–59^ and non-human primates^4,60^ suggests evolutionary conservation, an anatomically and molecularly defined A13 has not yet been identified in humans. This absence limits our understanding of both this dopaminergic cluster and the broader internal organization of the ZI in the human brain.

Here, we provide evidence for a human homologue of the A13 nucleus by integrating transcriptomics, histology, and ultra-high field (UHF) MRI. Following the definition established in rodent work, we refer to the A13 as a TH-positive nucleus situated in the medial ZI^47,51,52,57^. First, we identified a TH-enriched subregion using *post-mortem* human microarray data. Using *ex vivo* 15.2T MRI and histology from the same specimen, we independently validated TH expression and refined the spatial mapping boundaries of the A13. Next, we demonstrated that this TH-positive subregion exhibits elevated quantitative T_1_ relaxation and T_2_* values relative to the rostral and caudal ZI using *in vivo* 7T MRI data^2^. Lastly, we performed joint embedding analyses to define a principal axis of molecular and structural connectivity variation in the human ZI, providing evidence that the A13 likely represents a local molecular specialization Taken together, these findings provide converging evidence for a human A13 homologue and a rostral-caudal axis of ZI organization characterized by gradual transitions in molecular and anatomical connectivity, while presenting an approach integrating transcriptomics, histology, and neuroimaging to resolve fine-scale structures of the human brain. Understanding the human A13 also enables the translation of mechanistic insights from rodent models into humans, and informs the optimization of imaging contrasts within the deep brain.

## Results

### Evidence for a dopaminergic A13 subregion in the human ZI

Since the A13 is classically identified as a TH-rich region in the ZI^47,51,61^, we first examined whether such a subregion could be delineated from microarray data **(Figure 1)**. Gene expression associated with the ZI was extracted from the Allen Human Brain Atlas (AHBA, n=36 ZI-relevant samples from six donor brains, accessible at https://brain-map.org) using a validated atlas of the human ZI^2^. We performed principal component analysis (PCA) using a set of well-established cytoarchitectural gene markers that describe the ZI subregions^4,7,13,62^, with the first three principal components explaining 44.4%, 18.3%, and 16.7% of the variance respectively. Unsupervised clustering (k=5) produced five spatially coherent groupings within the ZI **(Figure 2a)**. When mapped back to the anatomical space, these clusters were located in the rostral, caudal, dorsal, and ventral regions within the ZI, along with a discrete cluster in the rostromedial region **(Figure 2a, Supplementary Fig. 1)**. Alternative solutions likewise resolved clusters along the rostral-caudal and dorsal-ventral axes, with a distinct cluster in the rostromedial region consistently parcellated across solutions (k=3-7 tested, **Supplementary Fig. 1**).

**Figure 1.**
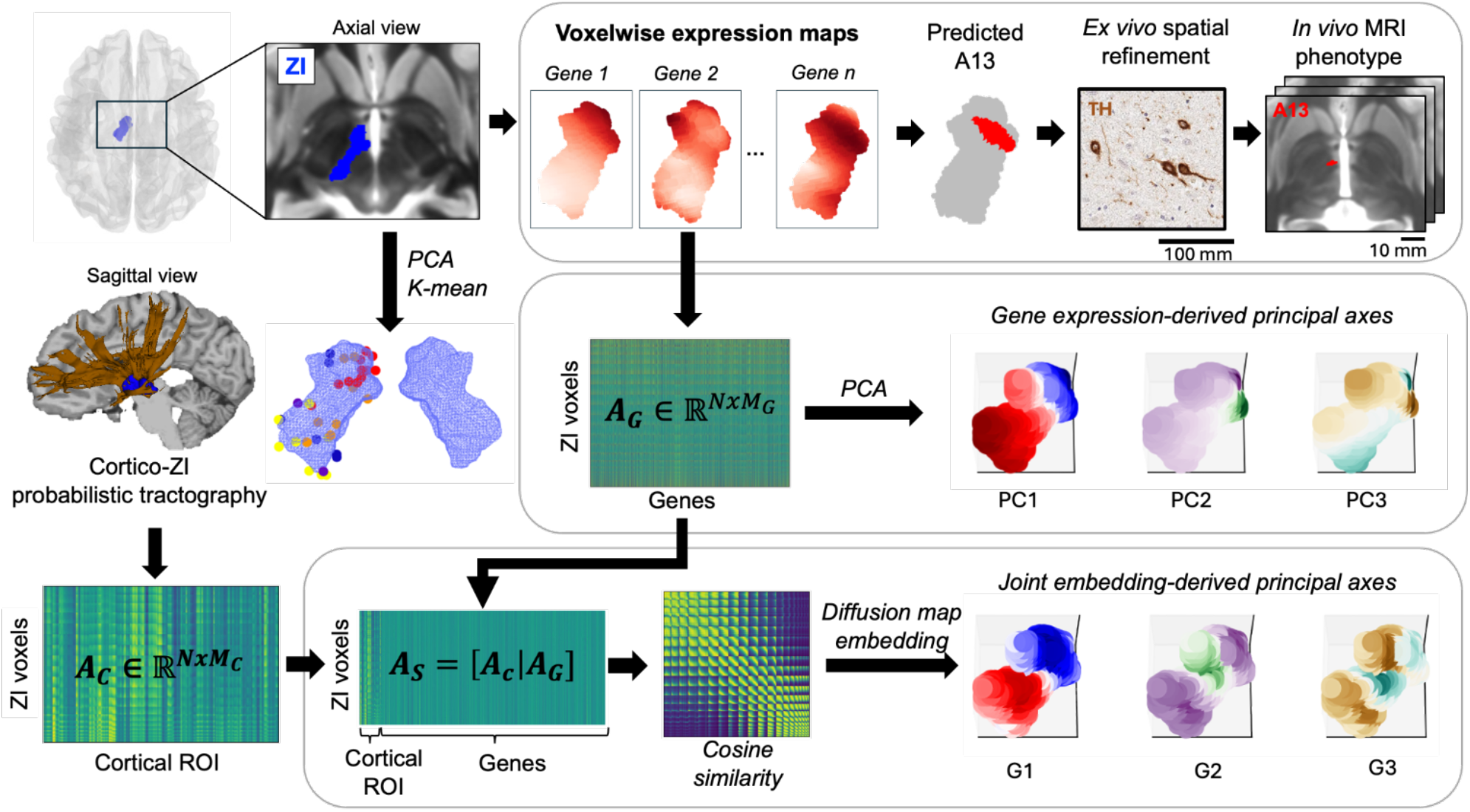
Multi-modal workflow for the spatial mapping of the human A13. Initial exploratory principal component analysis (PCA) and K-mean clustering of gene expression in the human zona incerta (ZI, axial view) identified a robust, tightly organized molecular cluster (red) in the rostromedial region, characterized by high tyrosine hydroxylase (TH) expression. *Top panel:* To improve spatial accuracy, we generated microarray-derived voxelwise expression maps and defined an A13 candidate mask by thresholding tyrosine hydroxylase (TH) expression at the top 15th percentile. We independently validated this mask using *ex vivo* 15.2T MRI and immunohistochemistry, confirming localized TH protein in the rostromedial ZI. Finally, the validated mask was applied to evaluate *in vivo* quantitative MRI phenotypes in healthy individuals (n=32). *Middle panel:* PCA of the voxelwise gene expression data (ZI voxel × gene matrix, A_G_) revealed principal axes of molecular variation in the human ZI, capturing a dominant rostral-caudal pattern (PC1). *Bottom panel:* Probabilistic tractography of cortical-incertal structural connectivity is represented by a ZI voxel × cortical ROI matrix (A_C_), and concatenated with the gene expression matrix A_G_ to produce a joint matrix (A_S_). Joint embedding (cosine similarity and diffusion maps) of the joint matrix revealed shared patterns of variation between the molecular and structural connectivity profile, identifying a dominant rostral-caudal gradient (G1).

**Figure 2.**
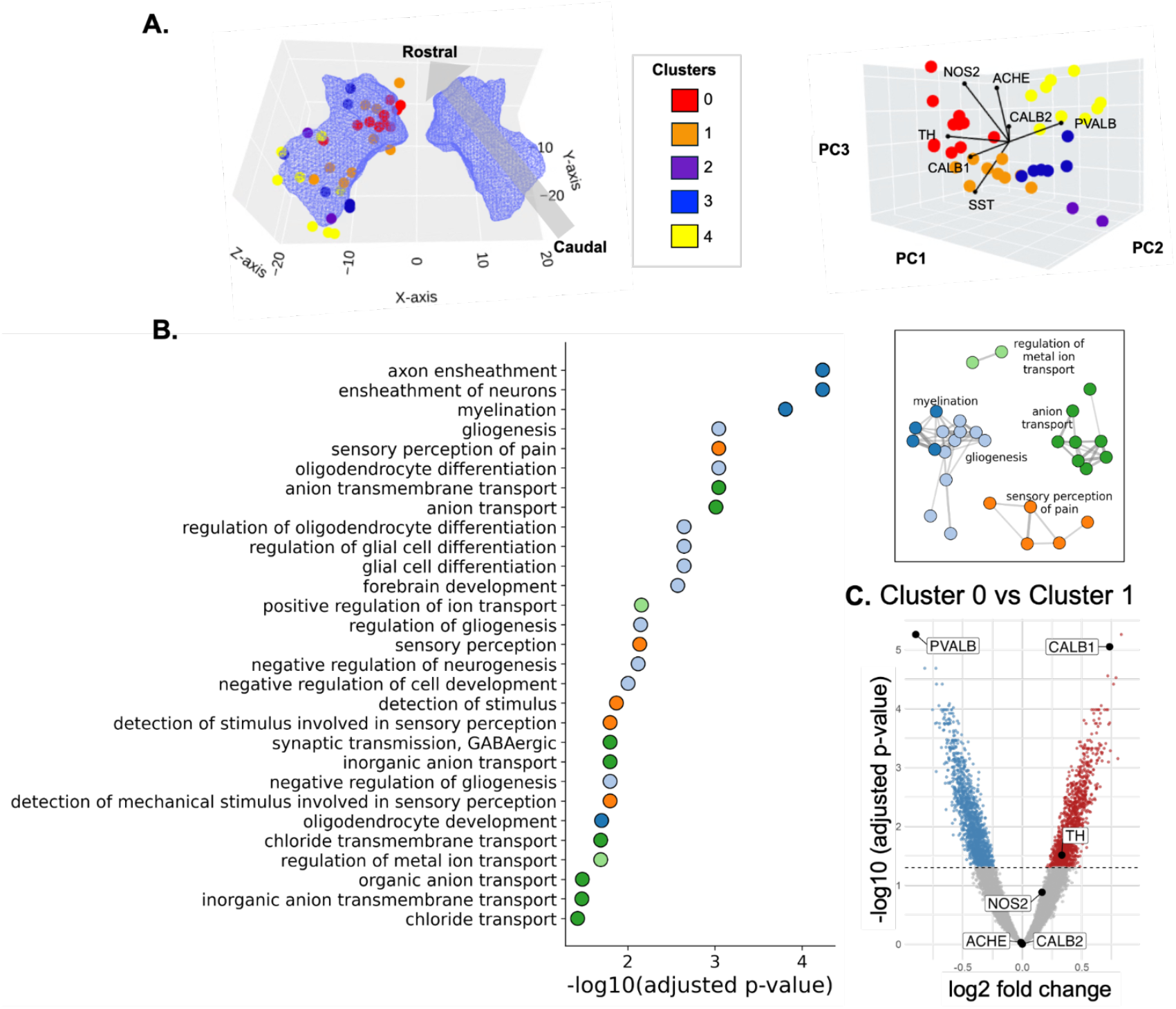
Transcriptomic-driven identification of a TH+ subregion in the rostromedial ZI. **A**. Principal component analysis (PCA) followed by unsupervised clustering reveal a tightly organized cluster in the rostromedial region of the human ZI (“cluster 0”, red). All identified clusters are visualized in MNI (*left panel*) and PC space (*right panel)*. Gene loadings are shown in PC space as represented by arrows. **B**. Over-representation analysis (ORA) identified significantly enriched pathways in cluster 0 relative to other ZI clusters. Enriched pathways were grouped into functional modules using the Louvain community detection algorithm (Networkx, v3.4.2, inset). Representative pathways include “sensory perception of pain” (GO:0019233, gene ratio=34/2453, adjusted p-value<0.0001) and “myelination (GO:0042552, gene ratio=129/2453, adjusted p-value<0.01)” within cluster 0. **C**. Volcano plot illustrates the molecular profile differences between cluster 0 and cluster 1, visualizing fold-change of gene expression (downregulated: blue; upregulated: red) and statistical significance (dotted line representing a threshold of −log10(0.05)). Cytoarchitectural gene markers in the ZI are highlighted, showing significantly upregulated TH expression in cluster 0 compared to cluster 1.

Differential gene expression between clusters revealed that the rostromedial region (“cluster 0”) was significantly enriched in TH expression relative to the rostral (“cluster 1”, adjusted p-value<0.001) and caudal region (“cluster 4”, adjusted p-value<0.001; **Figure 2c, Supplementary Fig. 2)**. We independently tested this observation through Marker Gene Profile^63^, a pipeline that enables the estimation of cell types in bulk tissue by summarizing gene expression data using known marker sets from RNA-seq and the literature. Dopaminergic markers were significantly enriched in the rostromedial “cluster 0” relative to the rostral “cluster 1” (adjusted p-value=0.042) and the caudal “cluster 4” (adjusted p-value=0.012; **Supplementary Fig. 2)**. As spatial coverage of sampling was relatively sparse in the central region of the ZI, we tested whether this affected our results by performing dimensionality reduction and unsupervised clustering on voxelwise expression maps within the ZI following established workflows^37,64^. Cluster and differential gene expression results were largely unaffected compared to the main findings (**Supplementary Fig. 3**). Taken together, these findings provide evidence for a molecularly distinct, TH-enriched cluster in the rostromedial region of the human ZI.

**Figure 3.**
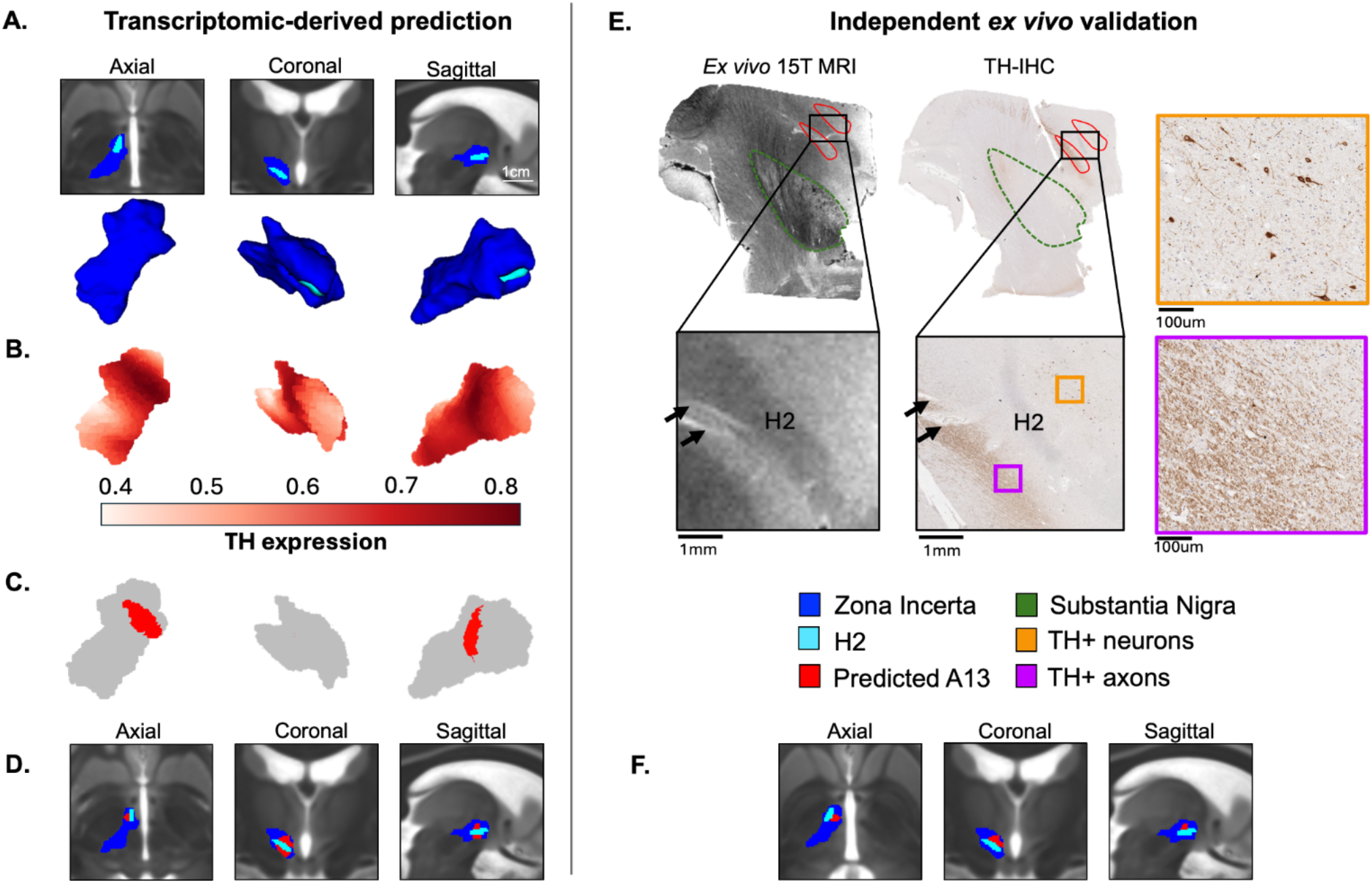
Spatial refinement of the putative A13 through ultra-high field *ex vivo* MRI and histology. **A**. The H2 fields of Forel (light blue) and the ZI (dark blue) as identified in the ZI 7T atlas^2^, visualized in MNI template space. **B**. Voxelwise tyrosine hydroxylase (TH) expression map of the ZI demonstrates the highest expression are located in the rostromedial region. **C**. Thresholding and binarization of the top 15th percentile of voxels containing the highest TH expression produced a candidate mask of the putative A13 (“predicted A13”, red). **D**. When visualized in MNI space, the predicted A13 mask occupies the rostromedial region of the ZI, superior and inferior to the H2 field of Forel. **E**. The predicted A13 mask was registered to the T1w volume of an *ex vivo* specimen, which guided serial sectioning at the predicted coordinates for immunohistochemistry (IHC). Using the same *ex vivo* sample, TH-IHC confirmed TH+ expression in neurons (orange inset) and axons (purple inset) in the predicted A13. H2 field of Forel is identified as an anatomical landmark. **F**. The putative A13 mask was refined based on histological pattern of TH+ neurons, restricting the A13 mask (red) to the region dorsal to the H2. Using previously calculated inverse transforms, we aligned the corrected mask back into MNI template space for visualization.

We next explored the functional associations of the TH-rich subregion through over-representation analysis (ORA). Cluster 0-specific genes were associated with functional terms including “sensory perception of pain” (GO:0019233, gene ratio=34/2453, adjusted p-value<0.0001), “forebrain development” (GO:0030900, gene ratio=93/2453, adjusted p-value<0.005), and “myelination (GO:0042552, gene ratio=129/2453, adjusted p-value<0.01)” (**Figure 2b, Supplementary Table 1)**, consistent with evidence of A13 function in rodents^47,65^.

### Multimodal mapping refines the spatial localization of the human A13

Previous studies have identified the A13 nucleus as a cluster of TH-positive neurons in the rodent^47,51,61^ and non-human primate brain^60^, and so we sought to delineate the putative A13 in humans in a similar manner. To this end, we generated a voxelwise TH expression map of the ZI and extracted the top 15^th^ percentile of voxels with the highest TH signal, yielding a discrete parcellation at a voxelwise resolution. This transcriptomic-driven parcellation localized the TH-enriched subregion to the rostromedial sector of the ZI [MNI centroid coordinates (mm): −6.2, −9.7, −6.4; volume: 73.6 mm^3^], surrounding the H2 (H2 field of Forel), a white matter tract known to traverse in the vicinity of the rZI^2,3,66^ (**Figure 3a-d)**.

To validate the transcriptomic predictions of the A13 boundaries, we conducted an independent, multimodal experiment integrating 15.2T MRI with histology using a cadaveric (*ex vivo*) specimen of the human subcortical brain, screened for absence of any underlying neurological disorder (see Methods). We used the 15.2T *ex vivo* image to delineate subcortical regions, and aligned it to the MNI template space. The predicted A13 parcellation was then registered into the native tissue space using the inverse transformation, which guided selective sectioning of the *ex vivo* specimen for immunohistochemistry (IHC) processing **(Figure 3c-f)**. This allowed us to bridge the spatial uncertainty inherent in the AHBA microarray dataset^67–69^ in IHC-based validation of TH+ expression within the ZI **(Supplementary Fig. 4)**. Furthermore, ultra-high magnetic field MRI enables increased signal-to-noise ratio (SNR) relative to conventional MRI, permitting direct visualization of subcortical structures which are not reliably resolved at lower field strengths ^2^ **(Figure 3a)**. When applied to *ex vivo* specimens, SNR is further boosted by prolonged scanning and absence of motion artifacts. Indeed, high resolution imaging of the dissected *ex vivo* specimen allowed the identification of gray matter regions of interest (i.e. ZI, STN, SNpc) and white matter tracts, including the H1 and H2 fields of Forel **(Figure 3e)**.

**Figure 4.**
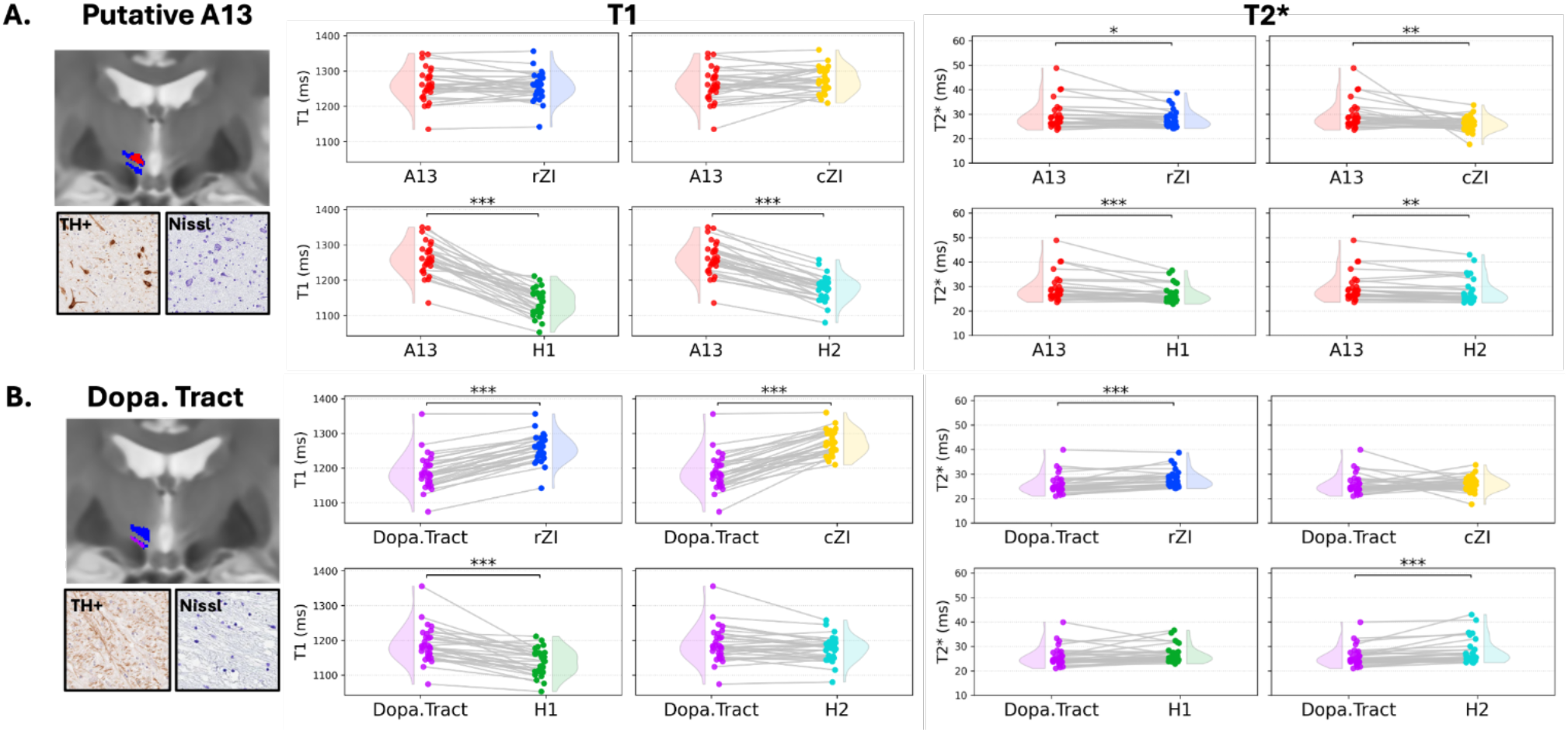
Delineation of the A13 region from adjacent structures through *in vivo* quantitative MRI. **A**. *Left panel:* The A13 (red) exhibits significantly longer T1 relaxation times compared to surrounding white matter tracts (H1, green; H2, cyan), and not significantly different from surrounding grey matter ZI subregions (rostral ZI: rZI, blue; caudal ZI: cZI, yellow). *Right panel:* The putative A13 also exhibits significant longer T2* values relative to both white matter tracts and adjacent ZI subregions. **B**. *Left panel:* The region occupied by TH+ axons (located ventral to the H2, purple) exhibits shorter T1 relaxation times compared to adjacent ZI subregions, but longer values when compared against nearby white matter tracts. *Right panel:* The dopaminergic tract exhibits shorter T2* values with rZI and H2, a relaxometry profile that is different from the TH+ neuron region.

By applying the described multimodal approach to naive tissue, we provided prospective validation of TH-expression and further spatial refinement of the transcriptomic boundaries of the predicted A13 parcellation. Importantly, this approach further revealed discrete cytoarchitectural landscape in the rostromedial ZI, distinguishing an area occupied by cell bodies of TH+ neurons **(Figure 3e**, orange inset) from an adjacent TH+ axon-rich region **(Figure 3e**, purple inset), both within the predicted putative A13 **(Figure 3d-e**, red region). TH+ neurons were restricted to the rostromedial region and not present within the central ZI (**Supplementary Fig. 5)**. The TH+ neuron cluster in the rostromedial ZI was located dorsal to the H2, while the TH+ axons are observed traversing ventral to the H2 and dorsal to the STN. Built on this new finding, we refined and restricted the boundaries of the putative A13 segmentation to include only regions dorsal to the H2, aligned with the histological patterns of TH+ neuron cell bodies [MNI centroid coordinates (mm): −6.7, −8.6, −4.6; volume: 34.4 mm^3^] **(Figure 3f)**. Together, these findings suggest increased spatial specificity for the localization of the A13 nucleus in the human brain.

**Figure 5.**
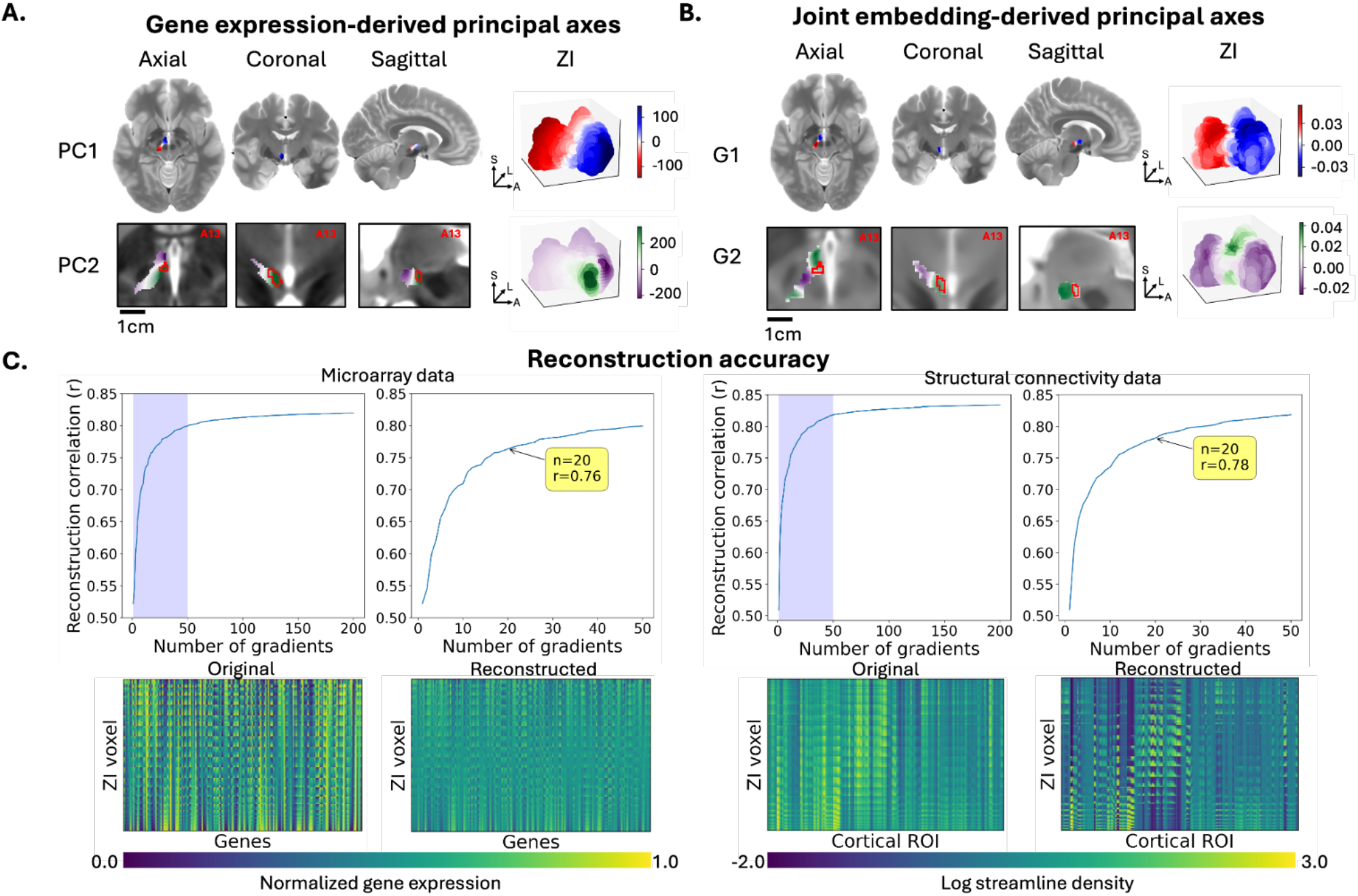
Principal axes of molecular and structural connectivity variation in the human ZI. **A**. PCA of the ZI gene-expression matrix identified two dominant principal axes, with PC1 and PC2 capturing variations along the rostral-caudal and medial-lateral axes, respectively. The positive-loading pole of the PC2 (*bottom panel*, green) overlapped with the A13 mask (red) in the human ZI (Dice coefficient: 0.55). Voxelwise PC score maps are presented in axial, coronal, and sagittal slice views. **B**. Joint embedding reveals shared pattern of variations of molecular and structural connectivity data along the rostral-caudal axis (G1), a pattern that is similar to PC1. The secondary joint gradient (G2) highlighted variation between the central ZI (*left panel*, green) and the rostral and caudal regions of the ZI. The A13 (red) does not align with any poles of the G2, unlike PC2, indicating that its separation from the rest of the ZI is not captured when structural connectivity data is incorporated. **C**. To ensure the joint gradients captured sufficient features from both modalities, the capacity to reconstruct the original molecular and structural connectivity matrices from each joint gradient was assessed. The original matrix from each modality was compared with the reconstructed matrix, where a high correlation coefficient represents higher amount of information captured. *Top panel:* The first 20 gradients captured majority of information from the microarray (Pearson’s r = 0.76) and structural connectivity data (Pearson’s r = 0.78). *Bottom panel:* Original and reconstructed matrices demonstrate preservation of information across ZI voxels for the microarray dataset (*left panel*, top 180 genes shown for visualization) and structural connectivity data (*right panel*, showing full 180 cortical ROI features)

### The human A13 homologue is distinctly identifiable in vivo at 7T

We next sought to identify whether an *in vivo* MRI signature exists for this *ex vivo-*validated TH+ subregion, which would provide the groundwork for future neuroimaging research. Quantitative T1 maps have been reported to be reliable and accurate for delineating the human ZI, as their intensity profile is similar to classic myelin-stained histological atlas^2,3^. Therefore, we compared the quantitative T1 values derived from 7T MP2RAGE images for the TH+ neuron subregion with surrounding structures in healthy participants (n=32; females=12; age: 46.2 ± 13.5 years^2^). The TH+ neuron subregion (1261.6 ± 46.8 ms) demonstrated significantly longer T1 relaxation times than the H1 (1137.2 ± 38.1 ms, adjusted p-value<0.001) and H2 (1176 ± 38.1 ms, adjusted p-value<0.001, **Figure 4a**). T1 values did not significantly differ between the TH+ neuron subregion and the rZI (1255.5 ± 39.8 ms, adjusted p-value=0.404) and the cZI (1273.2 ± 35.7 ms, adjusted p-value=0.162). Moreover, the TH+ neuron cluster could be distinguished from the TH+ axon region based on T1 relaxometry alone **(Supplementary Fig. 6)**. These results suggest the TH+ neuron subregion as a gray matter territory, sharing a common T1map signature with the broader ZI while remaining distinct from adjacent white matter fibres.

We next assessed T2* relaxometry as a complementary measure, given its sensitivity to iron content in subcortical dopaminergic nuclei such as the SNpc^70^. The TH+ neuron subregion (29.3 ± 5.6 ms) is significantly elevated in T2* values compared to the H1 (26.7 ± 3.5 ms, adjusted p-value<0.001), H2 (27.9 ± 5.1 ms, adjusted p-value<0.01), rZI (27.8 ± 3.6 ms, adjusted p-value<0.05) and cZI (25.9 ± 3.0 ms, adjusted p-value<0.01, **Figure 4a**). T2* is also significantly increased for the TH+ neuron cluster compared to other surrounding subcortical structures (including SN, STN, and the TH+ axon region, **Supplementary Fig. 6**).

The T1 and T2* values of the rostromedial ZI region occupied by TH+ axons were also evaluated. When compared across the same set of surrounding subcortical structures, the T1 values of TH+ axon region (1188 ± 51.6 ms) was significantly shorter compared to the rZI and cZI (adjusted p-value < 0.001), yet longer than H1 (adjusted p-value < 0.001, **Figure 4b**). The T2* values of the TH+ axon region were significantly shorter than that of the rZI and H2 (adjusted p-value < 0.001, **Figure 4b**), exhibiting a distinctly different relaxometry profile than the TH+ neuron region **(Figure 4, Supplementary Fig.6)**. Taken together, these quantitative MRI features indicate that the putative A13 containing TH+ neurons occupy a microstructurally distinct compartment within the ZI, with distinct T1 and T2* characteristics between gray matter ZI territories and adjacent white matter tracts.

### The human ZI is organized along major anatomical axes

Although the ZI has been classically categorized into discrete clusters based on neurochemical profiles^12,13,62^, little is known about how its molecular architecture may follow continuous axes of variation. Furthermore, while our findings indicate that the A13 is a distinct nucleus in the human ZI, it remains unclear how this subregion contributes to the broader gradient-based organization of the ZI.

To address these gaps, we first sought to investigate the organizational axes of the ZI by applying principal component analysis (PCA) to voxelwise gene expression maps associated with the ZI. The dominant axis of gene expression variation (PC1, 45.6% of variance) reveals a topographical gradient differentiating the rostral and caudal ZI **(Figure 5a)**. The PC1 score encoded position along the x and y axes of MNI template space, corresponding to the rostral-caudal dimensions (p-value < 0.001; ρ = 0.89, two-tailed CI=0.88-0.90).

The secondary principal component (PC2, accounting for 23.4% of total variance) captured variation along the medial-lateral axis. A compact and discrete region in the rostromedial ZI occupies its positive pole **(Figure 5a**, green region) while the rest of the ZI is captured by the transition zone and the negative pole (**Figure 5a**, purple region**; Supplementary Fig. 7**). PC2 is associated with the medial-lateral (ρ = 0.41, p < 0.0001, two-tailed CI = 0.45-0.38) and dorsal-ventral axes (ρ = 0.45, p < 0.0001, two-tailed CI = 0.48-0.41) in MNI standard space. The tertiary principal component (PC3, explained variance: 11.1%) varied along the dorsal-ventral axis of MNI standard space (ρ = 0.88, p-value < 0.0001, two-tailed CI = 0.89,-0.87, **Supplementary Fig. 7**), consistent with cytoarchitectural organization previously described in non-human primates^4,60^.

We observed that the primary (PC1) and tertiary (PC3) axes align with the established four-part subdivision of the ZI (rostral, caudal, ventral, and dorsal subregions^4,62^, **Figure 5a, Supplementary Fig.7**). In contrast, the second principal component (PC2) captures a distinct axis of variation along the medial-lateral axis: by overlaying the binary mask of the TH+ A13, we found that it overlapped with the positive-loading pole of the PC2, suggesting that the A13 may represent one of the primary drivers of molecular variance within the human ZI (Dice coefficient: 0.55; **Figure 5a**). Together, these results identify dominant patterns of variation in the ZI are aligned with major anatomical axes at the molecular level, within which the A13 emerges as a distinct subregion that may contribute to the secondary principal axis.

### The rostromedial ZI represents a molecular specialization along a conserved rostral-caudal axis

Having identified molecular-driven variations within the ZI, we next explored whether these principles are conserved across different biological scales, given that the broader ZI exhibits an organizational gradient in its cortical connections^5,10,11,71^. To achieve these goals, we performed probabilistic tractography between the ZI and neocortical targets, and conducted joint embedding between voxelwise gene expression data and cortico-incertal structural connectivity (see Methods). Joint embedding is a technique originally developed to compare connectivity organization across species by projecting data into a shared space and anchored to common features, such as homologous cortical regions between human and marmosets^44,45^. Here, we apply joint embedding to quantify shared patterns between molecular and anatomical connectivity data by mapping them into a common coordinate space. The resulting joint gradients are conceptually interpreted as shared patterns of variation between the investigated modalities^44,45^.

The primary joint gradient (G1) recapitulated a rostral-caudal axis of variation, consistent with the primary organizational axis (PC1) observed based on the microarray dataset (cosine similarity = 0.73; **Figure 5b, Supplementary Fig. 8**). This suggests that mesoscale connectivity and molecular expression share a common organizational scaffold. However, PC1 exhibited a rostromedial tilt relative to G1, suggesting that molecular variation may exist in this region that is not conserved in connectivity-based organizations. Moreover, the second joint gradient (G2) differs from the patterns of variation driven by microarray data (PC2). The G2 differentiated the central ZI apart from the rostral and caudal regions, rather than capturing a rostromedial subregion that overlaps with the A13, as illustrated by PC2 (**Figure 5a**,**b, Supplementary Fig.7,8)**. Together, these results indicate that the molecular and structural connectivity profiles in the human ZI vary along a rostral-caudal gradient. However, the A13 could not be separated from the broader ZI through this multimodal analysis **(Figure 5b, Supplementary Fig. 8)** or structural connectivity data alone (**Supplementary Fig. 10**, bottom panel). This discrepancy could reflect the possibility that the A13 represents a localized molecularly-defined subregion whose separation may not be captured by mesoscale connectivity.

### Reconstruction analyses of cross-modal joint gradients

To ensure the joint gradient captured sufficient information from both modalities, we evaluated how well the joint gradients could reconstruct each input matrix as described previously^38^. Reconstruction accuracy was quantified as the correlation between the original and reconstructed matrices across an increasing number of gradients, testing whether low-dimensional representations captured the dominant topology of each modality. Reconstruction accuracy of both modalities improved rapidly with first 20 gradients (reaching r ≈ 0.78 for structural connectivity and r ≈ 0.76 for gene expression matrix) and plateauing thereafter, with marginal gains up to 200 gradients (**Figure 5c**). G1 accounts for the largest proportion of information recovered (structural connectivity: r = 0.51, gene expression: r = 0.52), followed by G2 and G3 (structural connectivity: r = 0.10 and 0.15; gene expression: r = 0.04, r = 0.08).

We performed power spectral analysis to quantify the relative contribution of each joint gradient in reconstructing the original data^38^, revealing the gradients were supported by relatively balanced contributions between modalities rather than being solely dominated by either gene expression or structural connectivity **(Fig. 5c; Supplementary Fig. 8)**. Sensitivity analyses confirmed that the principal axis of variation is robust to alternative decomposition techniques (**Supplementary Fig. 9,10)**.

## Discussion

The A13 nucleus of the ZI has been well described in rodents^48,51,57^, yet no clear human homologue has been clearly identified. By combining multiple approaches, we provide the first detailed spatial mapping of the human A13 first in *ex vivo* data and then bridging this back to *in vivo* MRI. We further identify a rostral-caudal gradient conserved between biological scales in the ZI, evident at the molecular level and even when mesoscale structural connectivity is considered. However, while the A13 forms a discrete cluster and drives variance at the molecular level, its separation from the rest of the ZI is not maintained when structural connectivity is incorporated, suggesting the A13 may serve as a local molecular specialization in the human ZI.

Anatomically, the A13 nucleus is a catecholaminergic cell group in the mammalian dopaminergic system^48,49,59^, distinct from the paraventricular hypothalamic nucleus (A12) and the arcuate nucleus (A14)^72,73^. However, since the A13 is situated at the anatomical interface between the ZI and the hypothalamus, there remains debate as to which topographical region it belongs to. Consequently, it has been historically characterized using a variety of terms in the rodent literature such as the “medial ZI”, “rostral ZI”, “rostromedial ZI” or “incerto-hypothalamic region”^46,47,51^. In the present study, we adopted the anatomical localization from the rodent literature to identify the human A13 nucleus as a discrete TH+ cluster within the ZI, and we thus employ the use of an established parcellation^2^ to characterize this region.

We conducted independent histological experiments using spatial coordinates predicted by transcriptomics to refine our parcellation of the A13. This led to a spatial refinement of our localization of A13 as observations from high-resolution *ex vivo* MRI and histology indicate that the A13 region (defined by TH+ expressing neurons) is dorsal to the H2, STN, and furthermore that a bundle of TH+ axons courses through the rostromedial ZI **(Figure 3)**. While the dense convergence of these structures makes isolating specific tracts challenging, the anatomical profile of the dopaminergic tract could be compatible with the hypothesized trajectory of the superolateral medial forebrain bundle (slMFB^74,75^). Interestingly, both the A13 and slMFB are implicated in the modulation of valence and appetitive states^55,76,77^. Due to its potential in alleviating motivational deficits, the slMFB has been explored as a DBS target in treatment-resistant depression^77,78^ and obsessive compulsive disorder^79^ Its proximity supports the view that DBS in the vicinity of the rZI region may be a neuromodulatory target for neuropsychiatric indications^5,10^. Whether the therapeutic effect is due to slMFB or A13 stimulation, or some combined effect, remains an open question.

Together, our findings provide a framework for interpreting the potential functional significance of the putative A13. In rodent studies, the ZI as a whole modulates early circuit assembly^65^, while the A13 nucleus is more specifically implicated in sensorimotor modulation^50–52^, pain processing regulation^28,53^, and valence-based appetitive behaviours^55^. This offers a useful reference point for considering the functional properties of the human A13 homologue. The loss of *Arx*, a gene encoding for a transcriptional regulator, leads to failure of proper rZI dopaminergic neuron identity establishment in mice, subsequently leading to disrupted thalamocortical projections^65,80,81^. Our gene enrichment analysis aligns with these previous reports^65,80,81^: genes enriched in the putative A13 show significant association with “forebrain development”, “myelination”, and “glial cell differentiation”, pointing toward its role in shaping the cellular environment, supporting axon guidance, and the establishment of early subcortical-cortical connectivity. Enrichment for “sensory perception of pain” further implicates this region in nociceptive processing, in line with previous rodent studies^47,82^.

Due to its small size, direct *in vivo* visualization of the human ZI and its subregions has been challenging. However, certain tissue characteristics can be exploited when examined at higher magnetic field strengths. T1 dispersion (the dynamic range of T1 measurements) increases in a magnetic field-dependent manner^83^, which improves tissue contrast and distinguishes the ZI from surrounding grey and white matter regions^2^. T2* contrast similarly sharpens grey-white boundaries, as microstructural properties such as myelin and iron content modulate spin-spin relaxation and local susceptibility effects^84^. Building on prior work using UHF T1 mapping, we found that T1 relaxometry differentiated the putative A13 region from adjacent white matter tracts, and reproduced established T1 ranges for the rZI, cZI, and the fields of Forel^2^ (**Figure 4a**). Importantly, T2* values were significantly elevated in the identified TH+ region relative to neighboring structures, including the rZI, cZI, fields of Forel, and nearby dopaminergic pathways, further supporting the spatial specificity of these parcellations. Interestingly, the region occupied by the TH+ axons exhibited significantly elevated T1 relaxation values compared to H1, and reduced T2* relaxometry values than H2 **(Figure 4b)**. Although these relaxometry metrics are sensitive to myelin and iron content, the relaxation values represent a composite signal arising from a combination of factors, including water macromolecules, axonal orientation^85,86^, and iron oxidation states ^87^. Altogether, these findings indicate that the putative A13 exhibits a distinct profile identifiable using high-resolution MR relaxometry, as further evidence that this is a microstructurally distinct subregion.

Recent evidence supports the existence of topologically organized axes within the human brain^37,38,40–42^. A prime example is the sensorimotor-association axis, which is a fundamental gradient that bridges molecular, microstructural, and functional properties from primary sensory to high-order association cortices^41^. Consistent with this view, our findings show a rostral-caudal organizational axis identified independently using gene expression (PC1, **Figure 5a**) and structural connectivity data (published previously^11^), and on further analysis is conserved between these modalities (G1, **Figure 5b**). This indicates that an organizational scaffold along the rostral-caudal axis exists within the ZI across scales. Indeed, rodent studies demonstrate the GABAergic populations in rZI neurons preferentially influence fear and anxiety responses^17,20,34^, whereas glutamatergic cZI neurons modulate locomotion^35^. On the mesoscale connectivity level, primate tractography demonstrates an affective-motor dichotomy of rostral-caudal connectivity, with the rZI preferentially connecting with prefrontal regions and cZI with motor and somatosensory regions^5,10,11^. The identification of a principal rostral-caudal axis in the ZI provides further evidence that gradient-based architectures may represent a general principle of human brain organization.

Interestingly, we observed the A13 parcellation aligns with the rostromedial pole of the PC2, which is derived from molecular data **(Figure 5a)**, yet did not align with any gradients in joint embedding of transcriptomic and structural connectivity data **(Figure 5b, Supplementary Fig. 8)**. One plausible explanation is that the A13 represents a molecular specialization within the ZI by exhibiting a unique neurochemical profile, yet remains indistinguishable with the rest of the ZI when mesoscale cortico-incertal connectivity is examined. This suggests its function may rely more on molecular signalling rather than differences in mesoscale connectivity. Of course, we must also consider that the lack of structural separation may reflect the methodological limitation of tractography. Given the small size of the A13 nucleus and the complexity of crossing tracts in the surrounding region, its connectivity is incompletely resolved.

Though powerful, structural connectivity analyses also face additional methodological constraints: tractography cannot infer directionality and partial-volume effects in small nuclei may bias streamline estimates^88–90^. Imaging-transcriptomics using the AHBA dataset is further limited by relatively sparse gene expression sampling in its central region^91^ and is not without spatial bias. Nonetheless, the present study provides independent histological validation at the predicted coordinates to verify TH+ expression. This work offers a multimodal framework for examining the organizational axes of subcortical regions, providing a foundation for characterizing these nuclei across biological scales and offering pathways for clinical translation.

In summary, we have identified a molecularly distinct subregion in the human ZI consistent with the rodent A13, and demonstrate that it can be differentiated from surrounding structures using *in vivo* MRI. We further define a principal rostral-caudal axis in the adult human ZI, which is conserved when examined using molecular and structural connectivity approaches. Within this broader scaffold, the A13 subregion emerges as a local spatial and molecular specialization, highlighting how multimodal datasets can be leveraged across scales to enable a nuanced characterization of subcortical structures.

## Methods

### Human gene expression data

Voxelwise gene expression maps of the human ZI were obtained using the post-mortem microarray dataset from the Allen Human Brain Atlas (AHBA), consisting of 3702 samples from six donor brains^91^. Microarray samples within the human ZI coordinates^2^ (See *In vivo neuroimaging data and ZI mask* section) were extracted via the default parameters in the abagen toolbox^67^ as previously described^37,64^. As samples from the right hemisphere were only present in two subjects, only samples from the left ZI were included in analysis, followed by quality control through visual confirmation to ensure samples were within the ZI parcellations (raters=2, sample n=36). Discrete sample points passing quality control were used for downstream cluster analysis.

Microarray data sampled at discrete coordinates were interpolated across the volumetric space using ordinary kriging, a spatial model for estimating gene expression continuously across the subcortex^37,64^, resulting in voxelwise expression maps. Failed or unstable interpolation (dominated by zero-valued voxels) maps were excluded from downstream analysis, resulting in expression maps for 11,314 genes after quality control. To assess interpolation reliability within the ZI parcellation, leave-one-out cross-validation was performed by iteratively omitting each sample and comparing its predicted expression with the observed value **(Supplementary Fig. 4**). Specifically, for each ZI sample *i* (*i* = 1,2,…, *n*), we removed sample *i* and fit the same ordinary kriging model on the remaining *n* − 1 samples. The predicted expression was then compared to the observed expression at the location ŷ_i_. We summarized error with the root mean square error (RMSE), calculated as:

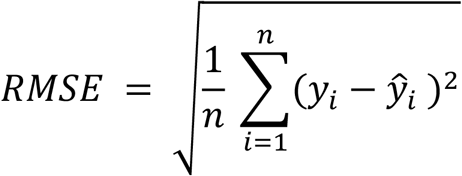

Voxelwise gene expressions are concatenated into a N-by-M_G_ matrix (1981 × 11314), where N represents ZI voxels in MNI152NLin6Asym space and M_G_ as genes. Continuous gene expression maps were used for downstream molecular gradient and joint embedding analyses.

### Molecular cluster analyses

Principal component analysis (PCA) was performed using the discrete microarray sample points within the ZI, using a curated gene set informed by rodent^7,13,62^, non-human primate^4,62,92^, and human^13,62^ literature of neurochemical markers for unique neuronal subpopulations in the ZI subregions: acetylcholinesterase (*ACHE*), calbindin 1 (*CALB1*), calbindin 2 (*CALB2*), nitric oxide synthase 2 (*NOS2*), parvalbumin (*PVALB*), somatostatin (*SST*), and tyrosine hydroxylase (*TH*). The top 5 principal components (PCs) captured a cumulative variance of 94%. Following PCA, K-means clustering was performed using a solution of k=5 based on previous cytoarchitectural knowledge of the human ZI^4^, with additional k-solutions tested (**Supplementary Fig. 1)**. Clusters were then visualized in MNI coordinate space overlaid with surface representation of the human ZI parcellation^2^.

Differential gene expression (DGE) between clusters were tested using the limma package on log-transformed expressions (v3.54.2). A design matrix encoding cluster membership for each gene was constructed, with the mean expression value of the given gene estimated for each group through linear models. Pairwise differences between clusters were tested for all genes using moderated t-test, followed by Benjamini-Hochberg false positive rate correction. Findings were tested in parallel through an independent pipeline^63^ **(Supplementary Fig. 2)**.

Significant genes surviving FDR corrections (threshold = 0.05) were mapped to Entrez IDs. Gene Ontology (GO) over-representation analysis was performed with clusterProfiler (enrichGO, clusterProfiler v4.6.2), with all genes identified in the ZI used as the background universe. The top 35 entries in the “Biological Processes” ontology were visualized through Networkx (Louvain algorithm, v3.4.2^93^) as a weighted graph, where enriched GO terms were represented as nodes. For a given pair of nodes, edges were defined by the Jaccard similarity coefficient, calculated as the ratio of shared DEGs between terms to the total number of unique DEGs associated with both terms. The Louvain community detection algorithm (default parameters, resolution = 1.0) to cluster pathways with overlapping gene sets, or similar biological functions, into cohesive modules.

### Ex vivo neuroimaging data

Brain specimens from a donor (female, age 75, no known neurological disorders) were obtained through the Western Body Bequeathal Program at Western University. Upon donation, the brain was fixed in situ by perfusion with 4% paraformaldehyde (PFA) through the carotid arteries, followed by fixation in 10% PFA for at least 14 days. The whole brain was dissected to extract the subcortex, and stored in 0.1% sodium azide-phosphate buffer solution at −4°C. The tissue was imaged overnight on a Bruker 15.2T scanner (Centre for Functional and Metabolic Mapping, CFMM) using a 3D fast low-angle shot (3D-FLASH) gradient-echo sequence with the following parameters: TR = 30 ms, TE = 8 ms, flip angle= 25°, matrix size = 268 × 358 × 224, bandwidth = 155 Hz/pixel, isotropic voxel size = 0.1 × 0.1 × 0.1 mm^3^, 4 averages.

Following imaging acquisition, the ex vivo T1w image was registered to the high-resolution 7T ZI template space^2^ through affine and non-linear transformations, with the inverse transformation used to propagate the transcriptomic-derived, predicted TH-rich parcellation into *ex vivo* subject space. Informed by the predicted coordinates, the specimen was cut into 1 cm blocks and paraffin-embedded to be serially sectioned at 4 μm intervals. All sections were stained using anti-Tyrosine Hydroxylase antibody (Milipore Sigma, cat# MAB318, lot# 4140356) and Nissl. Stained slides were mounted and digitalized at 20X resolution (Aperio Slide Scanner, Western Pathology Core). All procedures were complete as approved by the Western Research Ethics Board at Western University (REB# 123994), with written consent obtained from donors and families.

### Parcellation construction and refinement

Voxelwise tyrosine hydroxylase (TH) expression within the human ZI was first used to identify a transcriptomic-defined TH-rich region. Specifically, we extracted the top 15th percentile of voxels with the highest TH expression, yielding an initial transcriptomic-derived parcellation. This predicted TH-rich region was then compared with two independent anatomical references: (1) the rostral ZI (rZI) mask published previously^2^; and (2) TH-immunohistochemistry (TH-IHC) data digitized from serial histological sections. To generate a refined TH-rich parcellation, we retained only the voxels present in the intersection of all three sources: the predicted TH-rich region, the published rZI mask, and voxels exhibiting positive TH+ neuronal staining in the histology dataset.

### *In vivo* neuroimaging data and ZI segmentation

We leveraged an open-sourced dataset of 32 healthy participants (M = 20, F = 12; age 46.2 ± 13.5 years, range: 20-70 years; hereafter referred to as the “7T ZI” dataset)^2^. Structural and susceptibility-weighted images were acquired through 3D MP2RAGE and 3D ASPIRE sequences on a Siemens Magnetom 7T head-only scanner (Siemens Healthineers, Erlangen, Germany) at the Centre for Functional and Metabolic Mapping (CFMM, Western University). An 8-channel parallel transmit / 32-receive channel coil was used^94^. Detailed imaging and pre-processing steps to generate quantitative T1 maps were documented in previous literature^2^. For R2* map generation, raw multi-echo GRE data was reconstructed using an in-house singular-value decomposition (SVD) method based on Walsh et al to obtain a least-square magnetization estimate. Non-linear least-squares fitting of the complex multi-echo signal using the Levenberg-Marquardt algorithm was performed to estimate the R2* signal, voxelwise frequency, and complex amplitude simultaneously^95–98^. All procedures were performed in accordance to the HSREB office at Western University (REB# R-17-156).

We use the ZI segmentation from the 7T ZI dataset^2^ for both probabilistic tractography (see *in vivo diffusion-weighted neuroimaging data*) and transcriptomic analysis (see *Human gene expression data*). The probabilistic ZI mask was first thresholded at 50%, binarized, and subsequently dilated with a 3 voxel radius followed by registration into the MNI152NLin6Asym space. This ensured that the seeding region used in tractography was consistent with the mask used to extract the microarray data from the AHBA dataset.

### MRI phenotype of ZI subregions

For each participant, T1-weighted images previously registered to the 7T ZI template through affine and deformable registration^2^. To quantify MRI phenotype characteristics across ZI subregions, binary parcellations of ZI subregions (corrected TH-rich parcellation, rZI and cZI) and surrounding white matter tracts (H1 and H2 fields of Forel) were transformed back into subject native space through the inverse transforms^2^. Voxelwise values within each ROI were extracted from quantitative T1 maps and T2* maps, with mean values per ROI derived for each subject. These subject-level ROI measurements were assembled into matrices of shape ROI × subject for each quantitative map. Statistical comparisons between ROIs were performed using two-tailed paired Wilcoxon signed-rank tests, restricted to subjects with non-missing data in both ROIs being compared. P-values were adjusted for multiple comparisons using FDR.

### Molecular gradient analysis

We reduced the dimensionality of the gene expression matrix (1981 × 11315) through PCA after z-score normalization, with the top 10 PCs capturing a cumulative variance of 99%. Each PC score was mapped to the ZI parcellation in MNI152NLin6Asym space for visualization. Gene loadings were extracted as the PCA component weight matrix, with genes ranked by loading magnitude. To identify genes contributing to each PC, gene-set enrichment analysis was performed for the top 100 genes with positive and negative loading for each PC, querying Gene Ontology Biological Processes, KEGG, Reactome, and cell-type terms. To relate patterns captured in each PC with anatomical axes, the MNI coordinates of all voxels in the left-hemisphere mask were computed via affine transformation. For each PC, Spearman’s rho was calculated between voxelwise component scores and the x, y, and z-coordinate vectors, providing an estimate of how strongly each molecular gradient aligned with left-right, anterior-posterior, and inferior-superior axes.

### *In vivo* diffusion-weighted neuroimaging data

Structural connectivity between the cortex and ZI was assessed using minimally pre-processed 7T data from the Human Connectome Project (HCP) Young Adult study^99,100^ (n=169, M=68, F=101; aged 22-35). Diffusion-weighted images were acquired on a Siemens Magnetom 7T scanner using a 1.05 mm^3^ isotropic resolution. The acquisition protocol included a spin-echo EPI sequence (TR=7000 ms, TE=71.2 ms, FOV=210 × 210 mm^2^) across three diffusion-weighted shells (b-values = 1000, 2000, 3000 s/mm^2^) with 90 directions per shell. Additionally, 18 non-diffusion-weighted (b=0 s/mm^2^) images were acquired to facilitate distortion correction and provide a baseline signal. Full acquisition details are as described in the HCP1200 data release ^99,100^. We leveraged the minimally pre-processed HCP data, which includes correction for gradient nonlinearity, head motion, eddy currents, EPI distortion correction, and B0 intensity normalization^99^.

Probabilistic tractography was performed using FSL’s probtractx^101^, with detailed pre-processing steps as reported previously^11^. All subjects were processed with FreeSurfer (v5.3.0-HCP)^102^ as a part of the minimal preprocessed data release^99^. The ZI binary mask (dilated by 3 voxels and threshold at 50%) was first registered from the 7T ZI template space into the subject native space (0.7mm^3^ isotropic resolution), following the procedure. The HCP-MMP1.0 surface parcellation was projected into volumetric space using a ribbon-constrained label-to-volume mapping function and FreeSurfer-derived surfaces^103,104^. Using the ZI mask as seed ROI and cortical regions as target ROIs, connectivity was modelled with 5000 streamlines per ZI voxel, resulting in subject-level probability maps that quantified the number of streamlines reaching the neocortical targets. Connectivity maps were transformed to MNI152NLin6Asym space and assembled into an N×M_C_ connectivity matrix (1981 × 180), where N represents the ZI voxels and M_C_ as the cortical target regions.

### Joint embedding analyses

Gene expression data was represented as an N×M_G_ matrix, containing expression of 11314 genes across 1981 ZI voxels. Similarly, averaged SIFT2-weighted structural connectivity data across subjects were represented as an N×M_C_ matrix, containing connectivity between 1981 ZI voxels and 180 neocortical regions.

As the gene expression and connectivity data differed in the number of features, we performed data reduction using singular value decomposition (SVD) and z-score normalized to ensure the data are on a comparable scale. We concatenated the gene expression and connectivity matrices along the feature dimension to form a multimodal voxel-by-feature matrix (N × M, **Fig. 1**). To quantify voxel-wise similarity in this joint molecular-connectivity space, we computed cosine similarity between all pairs of feature vectors. For voxel i and j, cosine similarity is computed as:

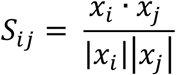

where *x* is the concatenated molecular-connectivity feature vectors. This resulted in an N × N affinity matrix A capturing how similar any two pairs of ZI voxels were in their combined molecular and connectivity profile.

We applied diffusion map embedding, a non-linear dimensionality reduction method, using the GradientMaps implementation in BrainSpace^105^. This approach identifies the principal axes of the data in a low dimensional representation, defined by the voxels’ multimodal similarity. The resulting gradients order voxels along a continuum based on their combined transcriptomic and structural profiles. For example, voxels characterized by a specific “signature”, such as high expression of Gene A and strong connectivity with Region B, are positioned at one extreme of the gradient; while voxels with the divergent profile (low expression of Gene A and minimal connectivity with Region B) are positioned at the opposite end. The optimal number of components was determined by inspecting an eigenvalue scree plot; following the elbow method, the first 20 gradients were retained for analysis^11,44,45^.

### Reconstruction analysis

We adapted the eigenmode-based reconstruction and power-spectrum analyses introduced by Pang et al., which were originally used to assess how cortical anatomy and functional MRI signals contribute to shared organizational patterns represented by eigenmodes (or “gradients”)^38^. In the present study, we generalize these analyses to quantify how molecular expression and structural connectivity contribute to the gradients derived through joint embedding. Each modality matrix *X* can be represented as *X ϵ* ℝ^*N*×*M*^, where N is the number of ZI voxels, and M is the feature (gene expression or structural connectivity). Matrix X was projected back onto the gradient basis *Ψ ϵ* ℝ^*N*×*n*^ for a given number of gradients *n*, where each column of *Ψ* contains gradients ordered by descending eigenvalue. We estimated the coefficient matrix *C*_*n*_ via the least-squares minimization:

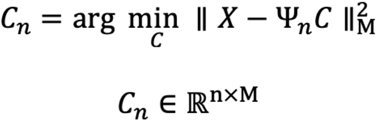

This produces gradient-by-feature matrix containing a set of coefficients (or ‘weights’) for the first *n* gradients. The reconstructed data for each modality is then obtained through:

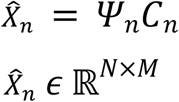

Reconstruction accuracy was quantified through Pearson’s r correlation between the original and reconstructed matrices:

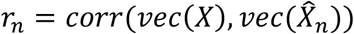

This analysis identifies the ability of the low-dimensional gradients at recapitulating the dominant spatial structure of each modality^38^. The number of gradients for reconstruction (*n* = 1…200) was chosen based on previous reports^38^.

### Power spectrum analysis

To estimate the relative contribution (“power”) of each modality to gradient, we computed a power spectrum from the coefficient matrix *C*_*n*_ = *C*_200_ *ϵ* ℝ^*G*×*F*^ for the first 200 gradients (*C*_200_) as described previously^38^. For modality X, power P of each gradient across features are computed as:

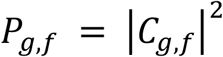

where g is a given gradient and f is a given feature in each modality. Power was then normalized within each feature, and averaged across features to obtain a single power score *P*_*g*_ per gradient:

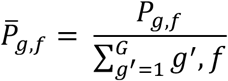

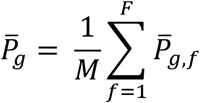

This procedure allowed for the identification of gradients that are driven by both molecular and structural connectivity.

### Sensitivity analyses

As AHBA microarray samples are acquired in donor space and subsequently registered to MNI space, MNI coordinates for the same tissue samples may vary depending on the registration algorithm employed. To assess whether these coordinate differences affected the predicted TH-rich parcellation, we repeated the procedures described in Parcellation Construction using three alternative coordinate sets:

1. Original MNI152NLin6Asym coordinates released by the Allen Brain Institute^91^;
2. Updated MNI152NLin6Asym coordinates released through the *alleninf* package^69^;
3. UpdatedMNI152NLin2009cSym coordinates released by the Cerebral Imaging Centre (CIC)^68^.

In addition, we conducted a series of sensitivity analyses based on alternative approaches used in prior gradient-mapping studies to assess whether alternative approaches of deriving shared spatial organization between modalities:

1. Performed SVD as joint decomposition^37^ instead of diffusion map embedding;
2. Performed diffusion map embedding independently to molecular and structural connectivity data;
3. Performed PCA independently to molecular and structural connectivity data.

For all alternative decompositions, the top components or gradients were chosen based on the elbow methods through scree plots, and retained to enable comparison of the dominant low-dimensional spatial axes across methods. PCA loadings, reconstruction and power spectrum analyses were performed to examine the relative contribution of each modality to the extracted components or gradients.

## Data Availability

All publicly available datasets analyzed in this study include: the post-mortem microarray data were obtained from the Allen Human Brain Atlas^90^ (https://brain-map.org); the structural and diffusion 7T MRI data for healthy adults from the Human Connectome Project Young Adult S1200 subjects data release (http://humanconnectome.org/study/hcp-young-adult)^99,100^; anatomical references and templates obtained from the 7T zona incerta atlas (https://osf.io/c8p5n/overview)^2^; and the MNI152 templates provided within FSL (v.5.0.11)^101^. The ex vivo dataset is available upon request.

## Acknowledgements

The authors would like to thank the individuals who donated their samples to the Western Body Bequeathal Program at Western University, without whom this research would not be possible. We also thank the members of the Western Body Bequeathal Program and the Pathology Core for their assistance, as well as the staff of the Centre of Functional and Metabolic Mapping (CFMM) for their technical expertise and support. This research was supported by the Canadian Institutes of Health Research (CIHR) and the Natural Sciences and Engineering Research Council of Canada (NSERC).

## Notes

### Competing Interest Statement

The authors have declared no competing interest.

