## Supplementary Figures for "An anatomically distinct dopaminergic cell population of the zona incerta as evidence for the human A13 nucleus"

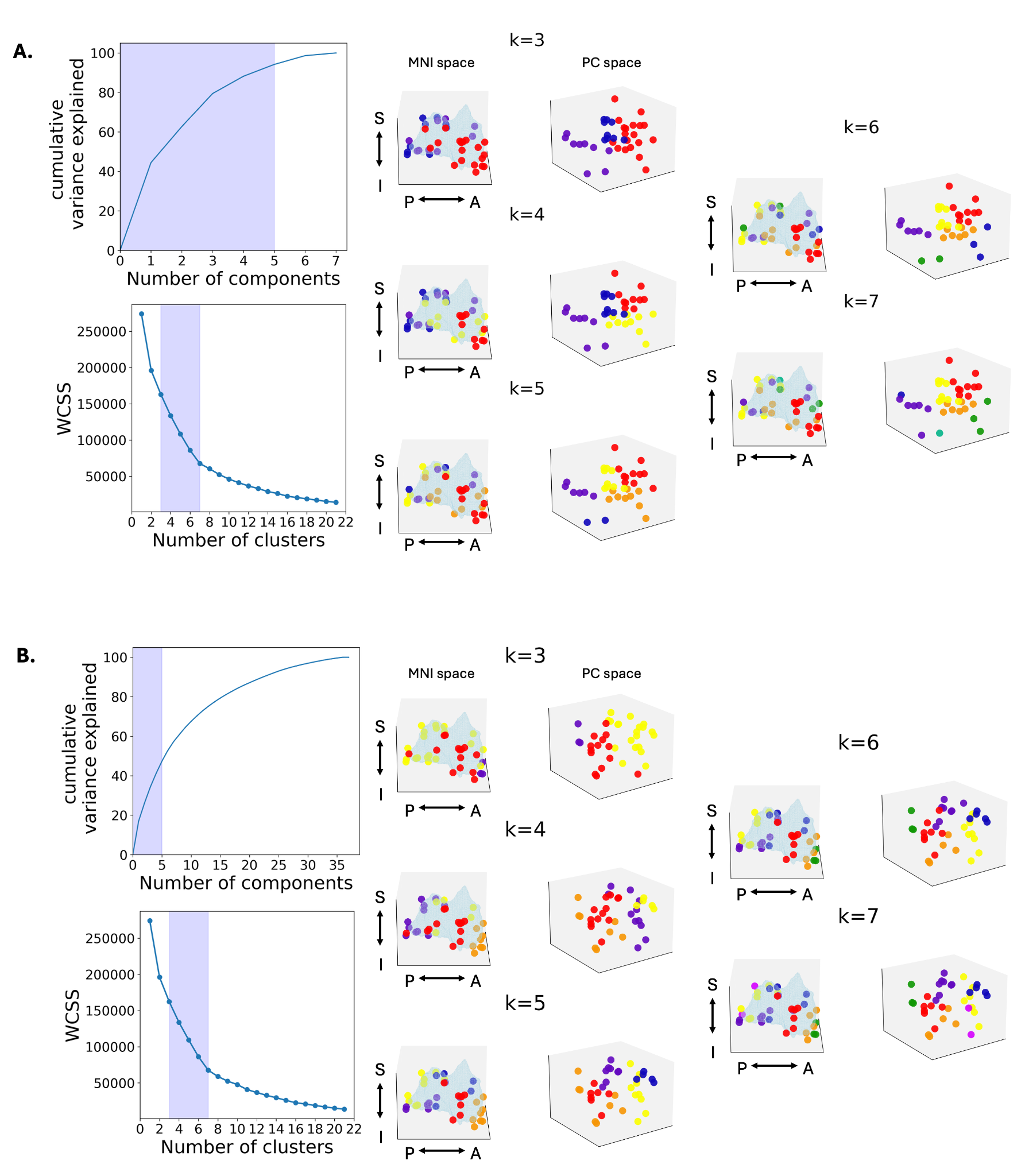

**Supplementary Figure 1. Additional K-mean solutions of ZI clusters.** **A.** PCA and k-means clustering (k = 3-7) were performed on a curated set of genes known to be relevant to ZI subregion neurochemistry (n=7 genes, See *Methods*). *Left panel:* Cumulative variance explained by principal components and within-cluster sum of squares (WCSS) across increasing cluster numbers. Shaded regions indicate the top 5 PCs and the range of k-mean solutions evaluated via the elbow method. *Right panel:* Spatial and molecular visualizations of k-means solutions in MNI space and PC space. Across solutions, K-mean reveals spatially coherent clusters within the ZI, with increasing k producing finer subdivision along the rostral-caudal axis. **B.** PCA and K-mean clustering (k = 3-7) was applied to the full set of all available genes within the AHBA microarray dataset (n=11,314 genes). *Left panel:* Cumulative variance and WCSS plots for the full gene set data, with k=3-7 selected through the elbow method. *Right panel:* Clustering solutions visualized in MNI and PC space, producing similar spatially coherent clusters.

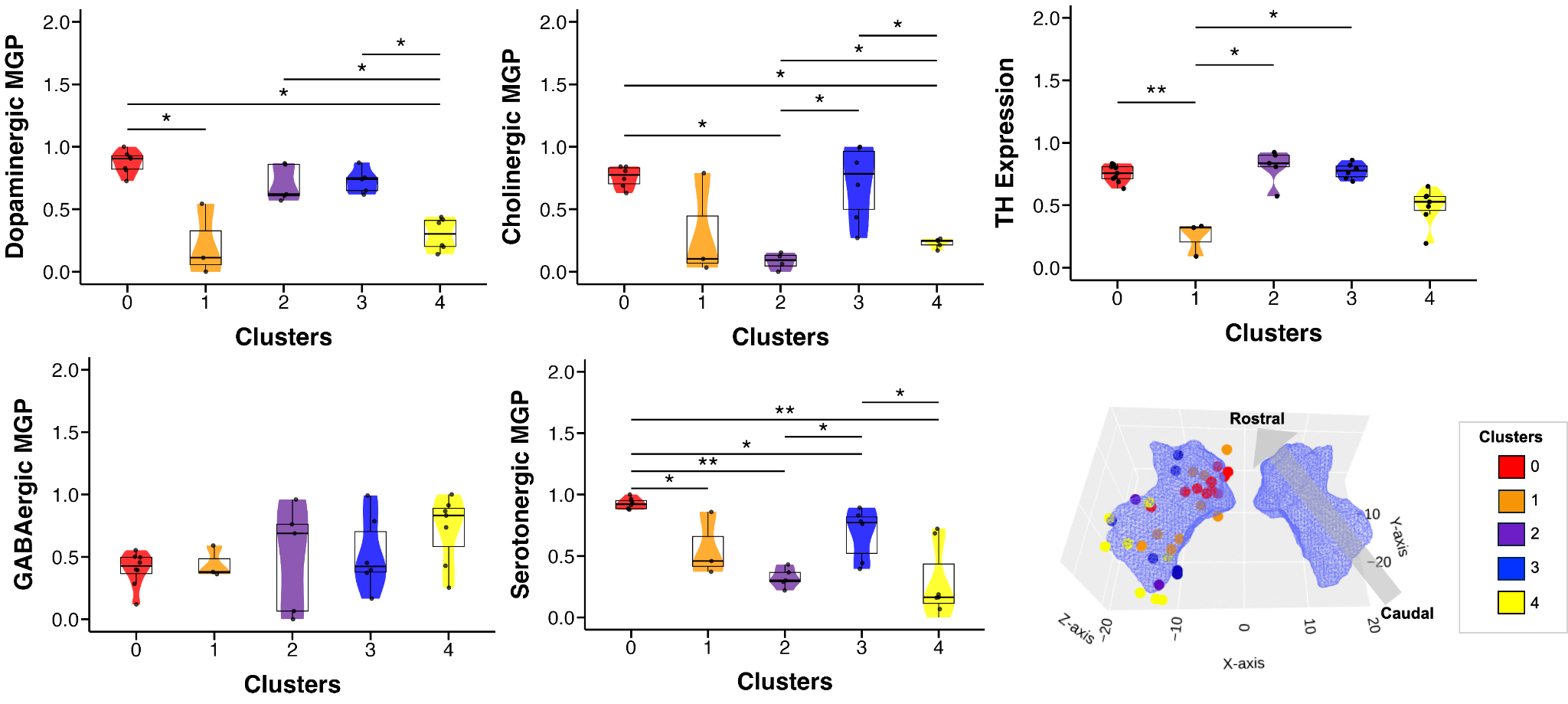

**Supplementary Figure 2. Elevated TH+ expression in the rostromedial ZI cluster.** *Left and middle panels:* Cluster-wise expression for normalized dopaminergic, cholinergic, GABAergic, and serotonergic marker gene panels (MGP) were tested through the MarkerGeneProfile^63^ package. Cluster 0 in the rostromedial ZI demonstrates a unique neurotransmitter profile, exhibiting significantly elevated dopaminergic, cholinergic, serotonergic marker gene expressions compared to Cluster 4, which is located in the caudal ZI. Clusters were defined by PCA followed by K-mean analysis on the AHBA microarray samples localized to the ZI parcellation surviving manual quality inspection. *Right panel:* Log-normalized differential TH expression across clusters recapitulates the expression pattern as demonstrated by dopaminergic MGP, showing significantly elevated TH expression in Cluster 0 of the rostromedial ZI compared to other subregions. Statistical comparisons were calculated via Limma empirical Bayes moderated t-tests with Benjamini-Hochberg correction (* adjusted p-value < 0.05, ** adjusted p-value < 0.01).

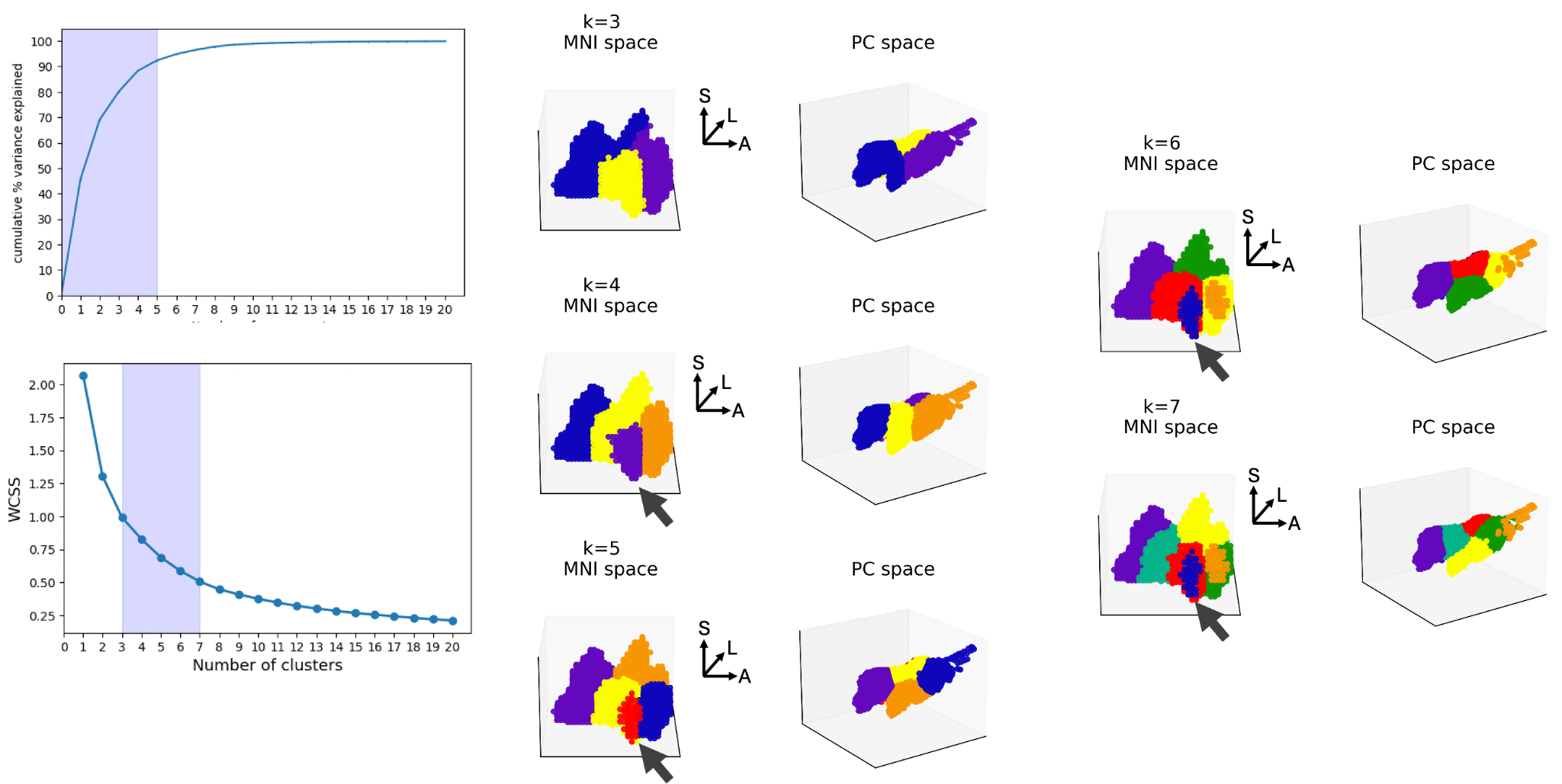

**Supplementary Figure 3. Additional K-mean solutions of ZI clusters using voxelwise expression maps.** *Left panel:* Cumulative variance explained by principal components and within-cluster sum of squares (WCSS) across increasing numbers of clusters. Shaded regions indicate the variance explained by the top 5 PCs and k-mean solutions tested based on the elbow method. *Right panels***:** K-means clustering solutions (k = 3–7), visualized in MNI space (left column) and in the corresponding PC space (right column). Across solutions, clustering consistently highlight a rostromedial region (arrow) and produced spatially coherent groupings within the ZI, with increasing k producing finer subdivision along the rostral-caudal axis.

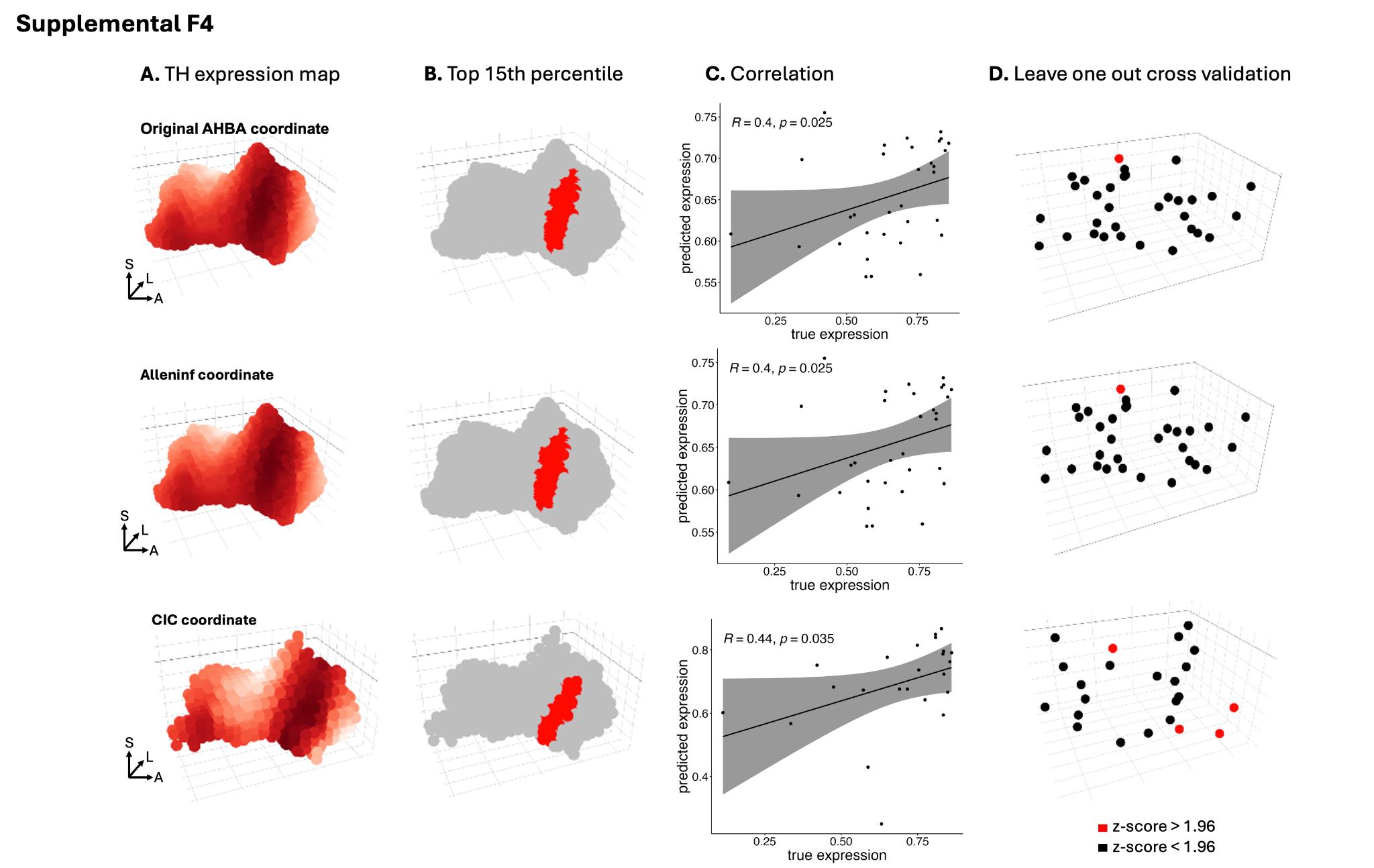

**Supplementary Figure 4. Robustness of TH expression-map based delineation of the putative A13 across coordinate systems. A.** Voxelwise TH expression map projected in the template space using the original coordinate sets (provided by the Allen Institute^91^), AllenInf coordinates (provided by Gorgolewski et al.^69^), and the Cerebral Imaging Centre (CIC) coordinate set^70^. Color intensity reflects normalized TH expression. **B.** Binary masks derived from the top 15th percentile of TH-expressing voxels in each coordinate system, highlighting a consistent rostromedial subregion within the ZI. **C)** Correlation between predicted and observed TH expression values across samples for each coordinate system, demonstrating significant positive associations. Shaded regions indicate confidence intervals of the regression fit. **D)** Leave-one-out cross-validation analyses showing stability of the predicted TH-enriched region across samples. Red points denote significant z-scores (> 1.96), black points denote non-significant values, with samples with high errors locating near the edge of ZI parcellation.

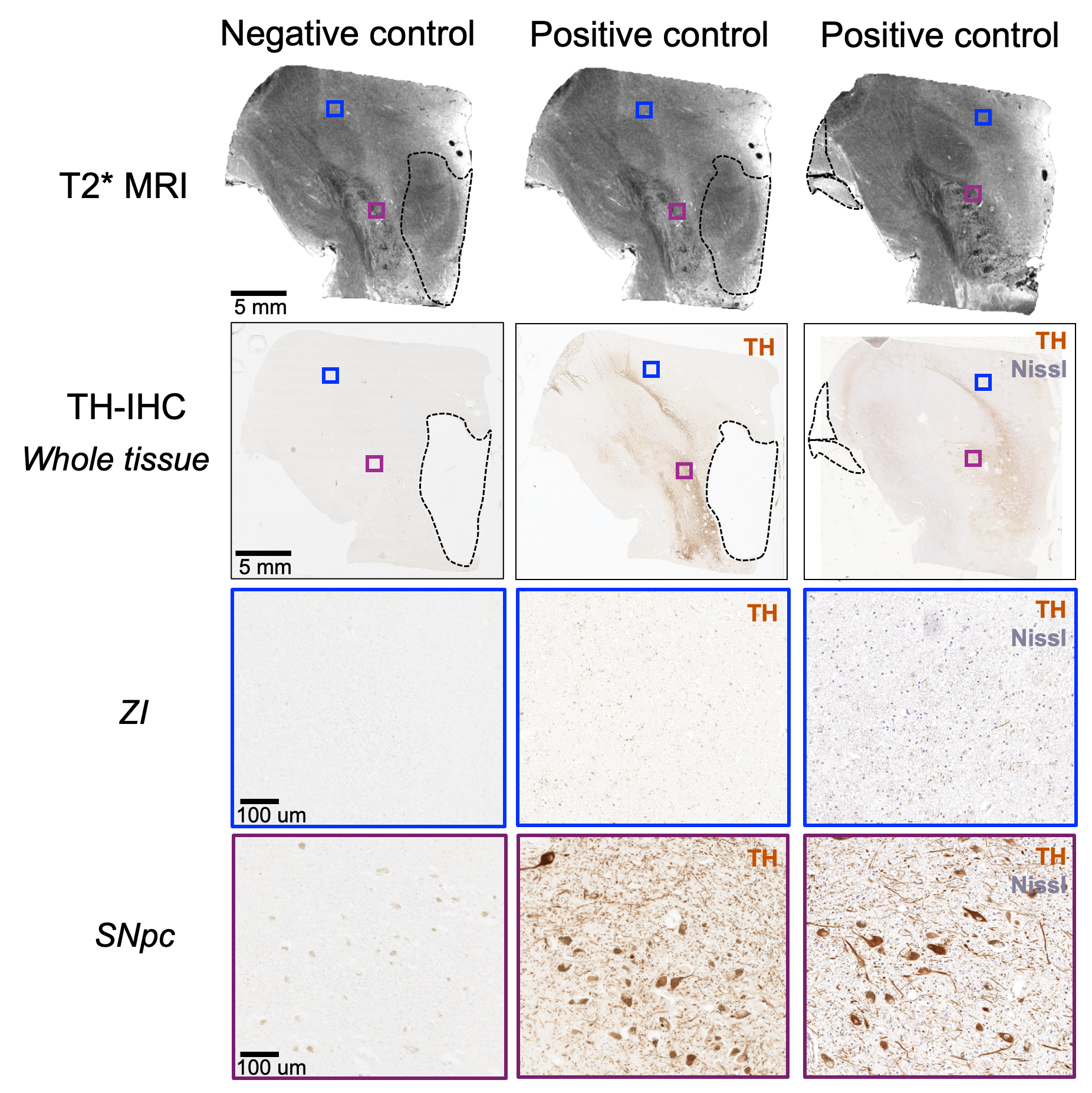

**Supplementary Figure 5. Immunohistochemistry validation of tyrosine hydroxylase expression in an independent *ex vivo* specimen.** Corresponding slices of the T2* MRI and TH-IHC from an independent *ex vivo* specimen shown in coronal view, with central region of the ZI (blue) and the SNpc (purple) insets highlighted. Negative control (no TH primary antibody or Nissl staining) is shown along with two positive controls (*middle panel:* TH primary antibody treatment; *right panel:* TH primary antibody treatment, counterstained with Nissl) from the same batch, showing specificity of the primary antibody. No positive neuron staining was observed in the central ZI, in line with previous studies confirming TH+ expression restricted to the rostral ZI. Strong TH+ signals in neurons are observed in the positive controls of SNpc. *ZI, zona incerta; SNpc, substantia nigra pars compacta*

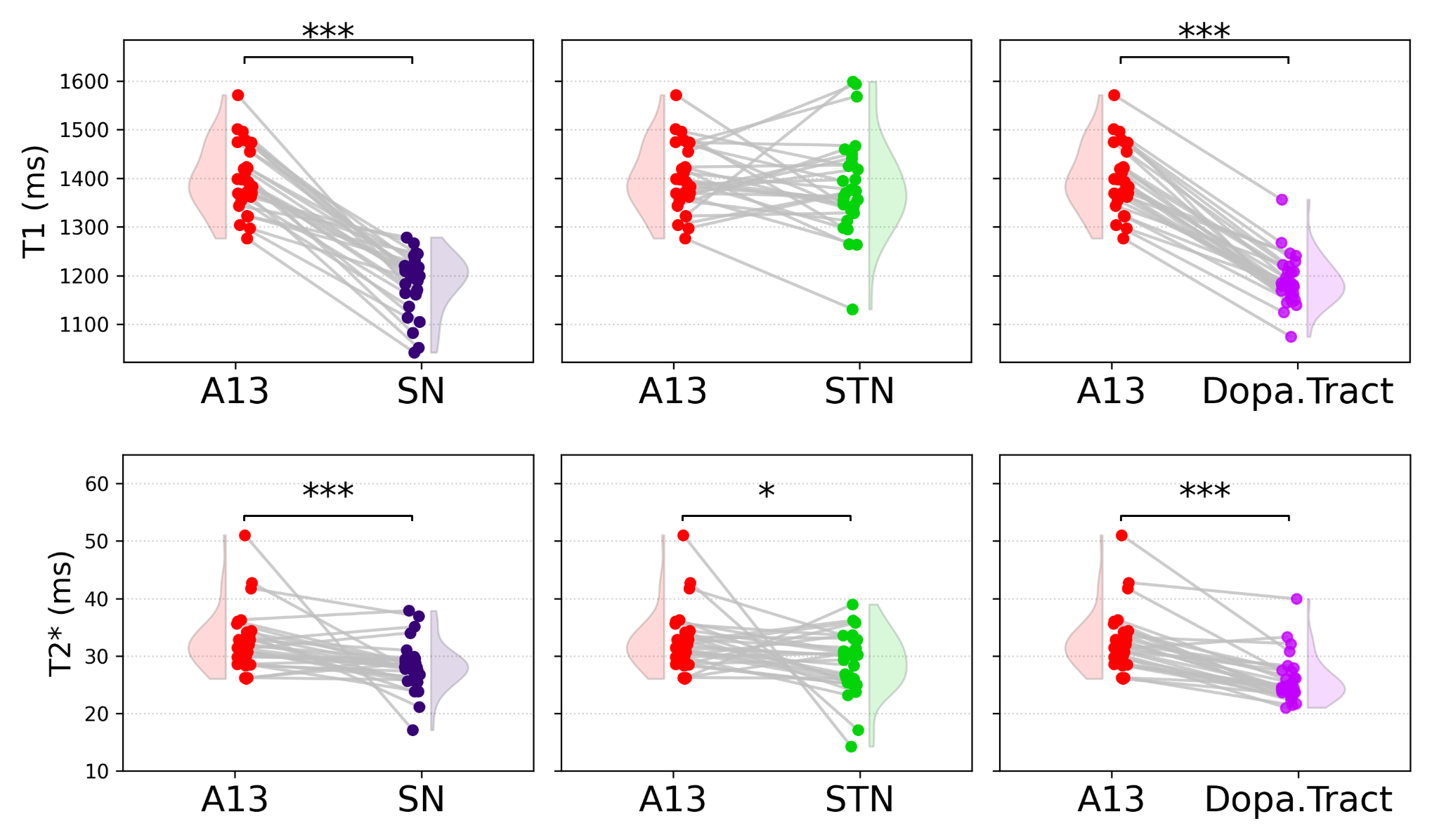

**Supplementary Figure 6. Quantitative MRI comparisons between the A13 and neighboring subcortical regions.** *Top row:* T1 relaxation time (ms). *Bottom row*: T2* (ms). Paired comparisons are shown between the putative A13 and substantia nigra (SN), subthalamic nucleus (STN), and the dopaminergic tract (Dopa.Tract) traversing inferior to the H2 Field of Forel in healthy individuals (n=32, Wilcoxon matched pairs signed rank test, *p < 0.05, ***p < 0.001; ns, not significant).

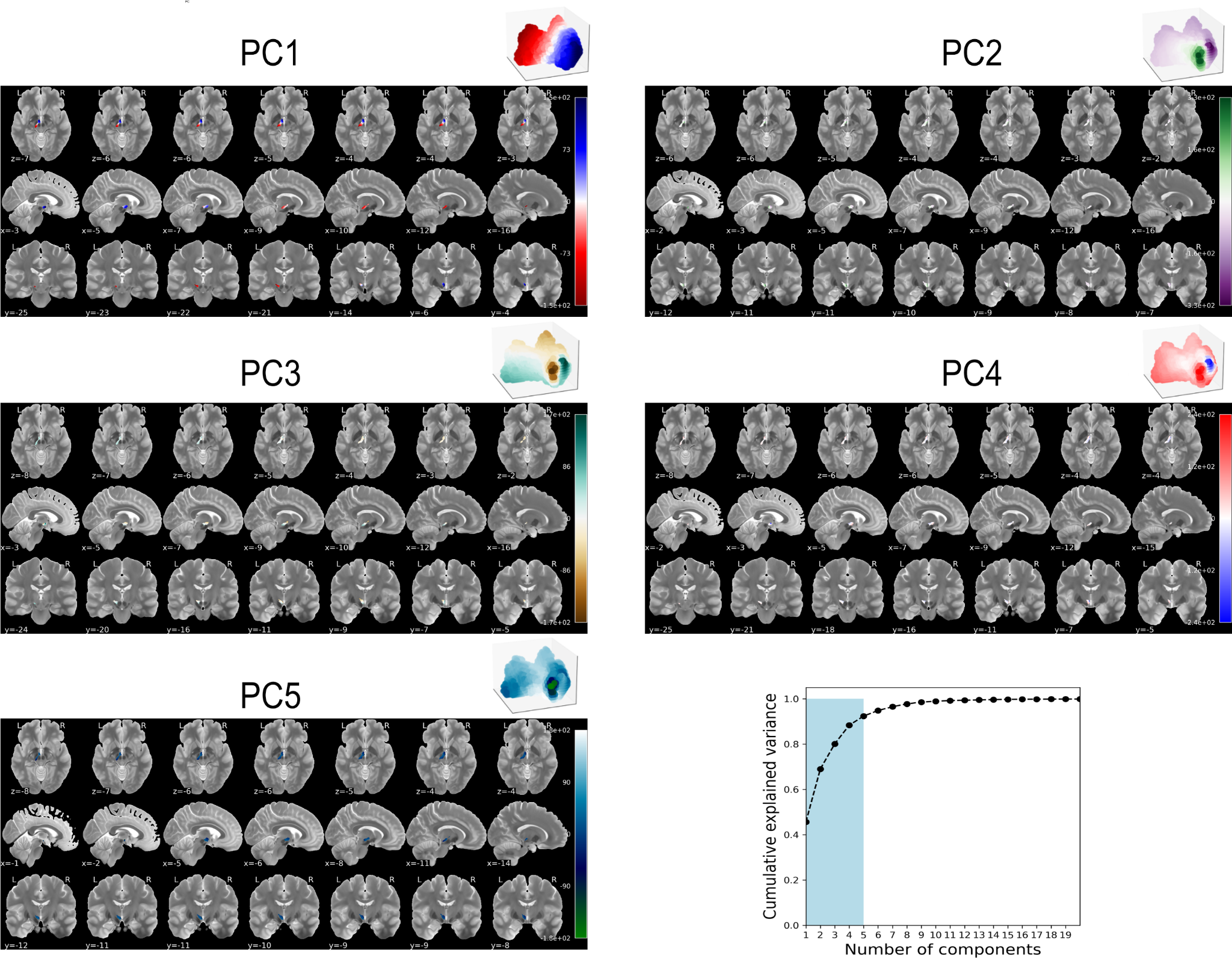

**Supplementary Figure 7. Visualization of principal components from microarray data.** Voxelwise visualization of the first five principal components (PC1-PC5) derived from the ZI gene-expression matrix are shown in axial, sagittal, and coronal views in MNI space. PC1 captures a dominant rostral-caudal molecular gradient, while subsequent components (PC2-PC5) represent orthogonal axes reflecting finer-scale spatial variation within the ZI. Cumulative variance explained as a function of the number of retained components is plotted with shaded region indicating the selected dimensional range used for downstream analyses.

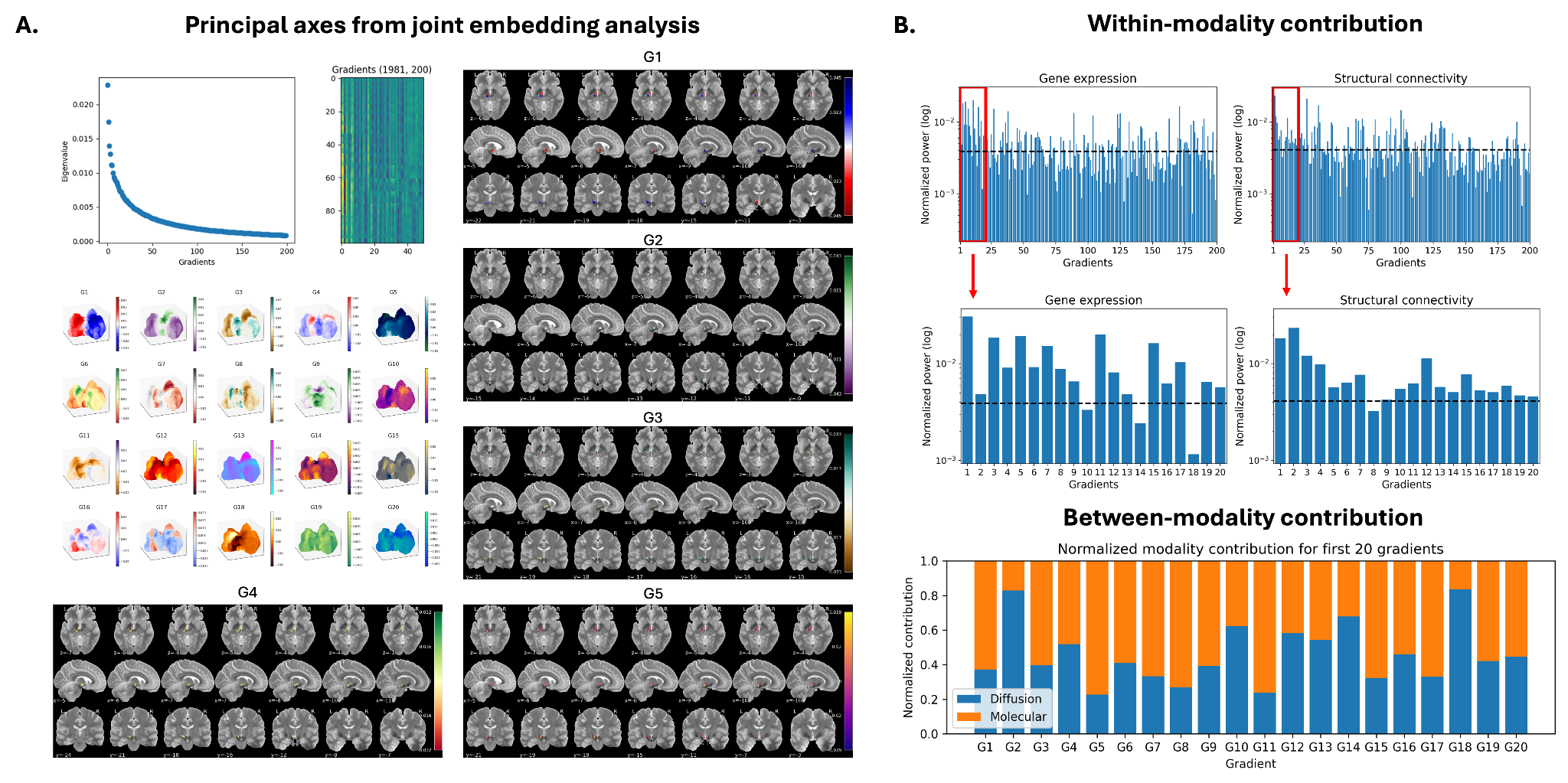

**Supplementary Figure 8. Joint embedding analysis of gene-expression and structural connectivity data. A.** Principal axes from joint embedding captured molecular and structural connectivity variation within the human ZI. *Top:* Meaningful number of gradients were identified as the inflection point (Gradient = 20) as identified by plotting eigenvalues as a function of gradient number. *Middle:* Voxelwise maps of the first 20 joint gradients (G1–G20) are shown in axial, sagittal, and coronal views in MNI space. G1 reflects a dominant rostral-caudal organizational axis, while subsequent gradients capture finer-scale spatial variation within the ZI. **B.** Power spectrum analysis across the top 200 gradients quantifying the contribution of each joint gradient to the original gene-expression and structural connectivity matrices (within-modality contribution, red box highlights the top 20 gradients), and; **C.**  the relative contribution of molecular and structural connectivity data to each joint gradient (between-modality contribution). Results indicate relatively balanced contributions across modalities, demonstrating that the joint gradients are not driven disproportionately by either molecular or connectivity matrices.

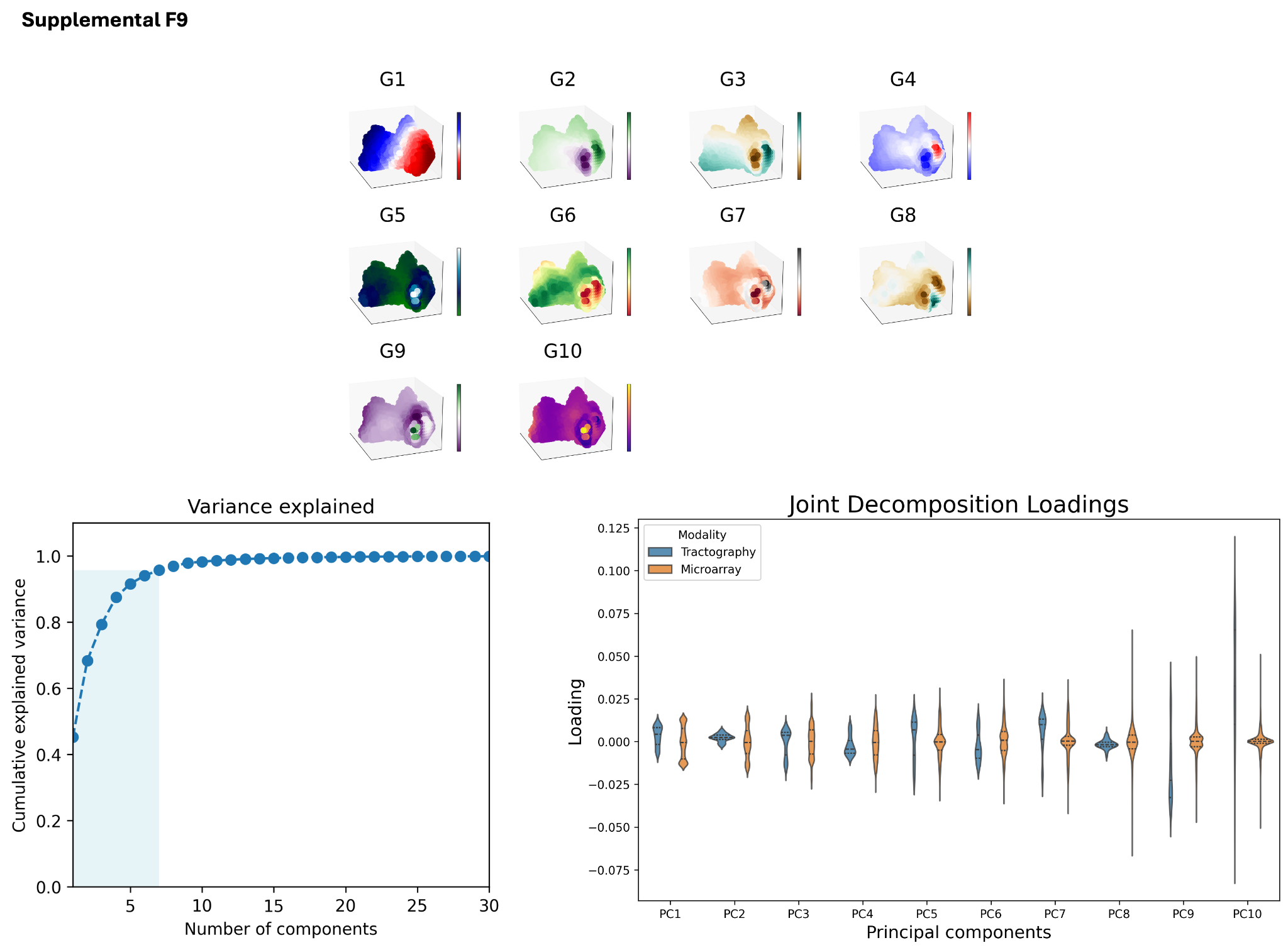

**Supplemental Figure 9. Cross-modality principal axes as revealed through joint decomposition.** Microarray and structural connectivity data were z-score normalized, and decomposed through SVD as described previously^43^. *Top:* Gene-expression and cortico-ZI structural connectivity matrices were concatenated together with dimensionality reduced through PCA, showing that a rostral-caudal gradient was preserved, yet molecular data dominants the joint gradients as visualized through PC loadings.

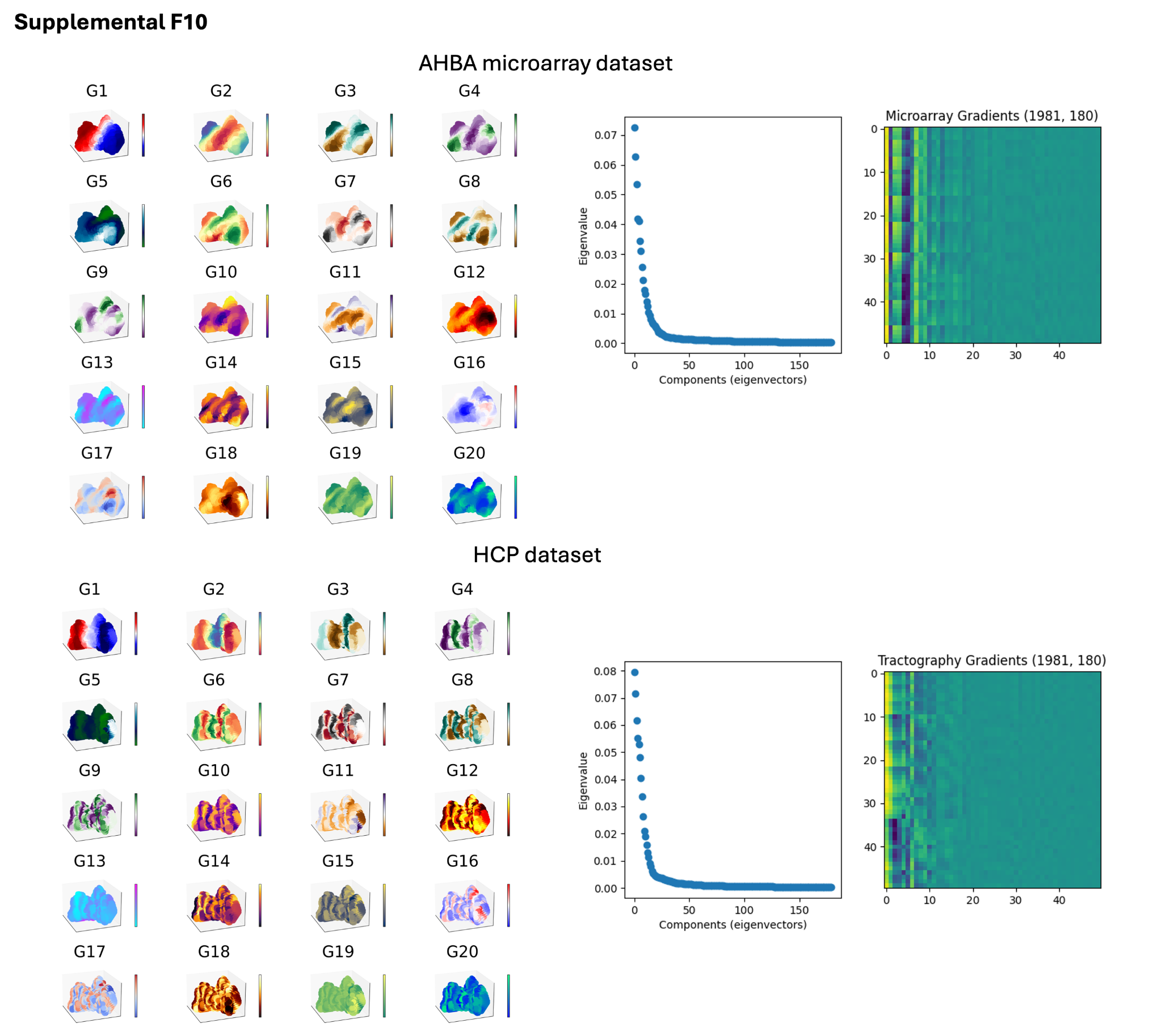

**Supplemental Figure 10. Diffusion map embedding of gene expression data and structural connectivity data in the human ZI.** Diffusion map embedding of gene-expression matrix (top) and structural connectivity matrix (bottom) produced internal organizational gradients (G1-G20) in the human ZI, showing that the rostral-caudal gradient (G1) was preserved.
